# The YΦ Motif Defines the Structure-Activity Relationships of Human 20S Proteasome Activators

**DOI:** 10.1101/2021.04.28.441858

**Authors:** Kwadwo A. Opoku-Nsiah, Andres H. de la Pena, Sarah K. Williams, Nikita Chopra, Andrej Sali, Gabriel C. Lander, Jason E. Gestwicki

## Abstract

The 20S proteasome (20S) facilitates turnover of most eukaryotic proteins. Substrate entry into the 20S first requires opening of gating loops through binding of HbYX motifs that are present at the C-termini of certain proteasome activators (PAs). The HbYX motif has been predominantly characterized in the archaeal 20S, whereas little is known about the sequence preferences of the human 20S (*h*20S). Here, we synthesized and screened ∼120 HbYX-like peptides, revealing unexpected differences from the archaeal system and defining the *h*20S recognition sequence as the Y-F/Y (YΦ) motif. To gain further insight, we created a functional chimera of the optimized sequence, NLSYYT, fused to the model activator, PA26^E102A^.A cryo-EM structure of PA26^E102A^-*h*20S identified key interactions, including non-canonical contacts and gate-opening mechanisms. Finally, we demonstrated that the YΦ sequence preferences are tuned by valency, allowing multivalent PAs to sample greater sequence space. These results expand the model for termini-mediated gating and provide a template for the design of *h*20S activators.

---

The proteasome is a critical regulator of protein homeostasis that degrades ∼90% of all eukaryotic proteins^1^. The enzymatic activity of this system is carried out by the 20S proteasome (20S), a cylindrical, complex composed of four stacked rings enclosing an axial channel. To be degraded, potential substrates must first diffuse through a pore at the center of the distal α-rings before encountering the peptidase sites within the inner β-rings. N-terminal extensions of the α-subunits “gate” entry into the pore, limiting the degradation of bystander proteins^2^. In turn, this barrier creates a key regulatory role for the proteasome activators (PAs)^3–5^, large particles that bind the 20S, open the gates, and facilitate substrate selection and entry. One evolutionarily conserved^6–8^ way that PAs achieve this goal is by using a tripeptide motif at their extreme C-termini, which is characterized by a hydrophobic amino acid, followed by a tyrosine and then any amino acid (HbYX). HbYX motifs open the gates by docking into pockets located between adjacent α-subunits of the 20S (termed α-pockets) (**Fig. 1a**)^9–11^. Unlocking the details of HbYX recognition at this protein-protein interaction (PPI) will deepen our understanding of the gate opening mechanism and enable novel strategies to regulate protein degradation in cells.

**Figure 1.**
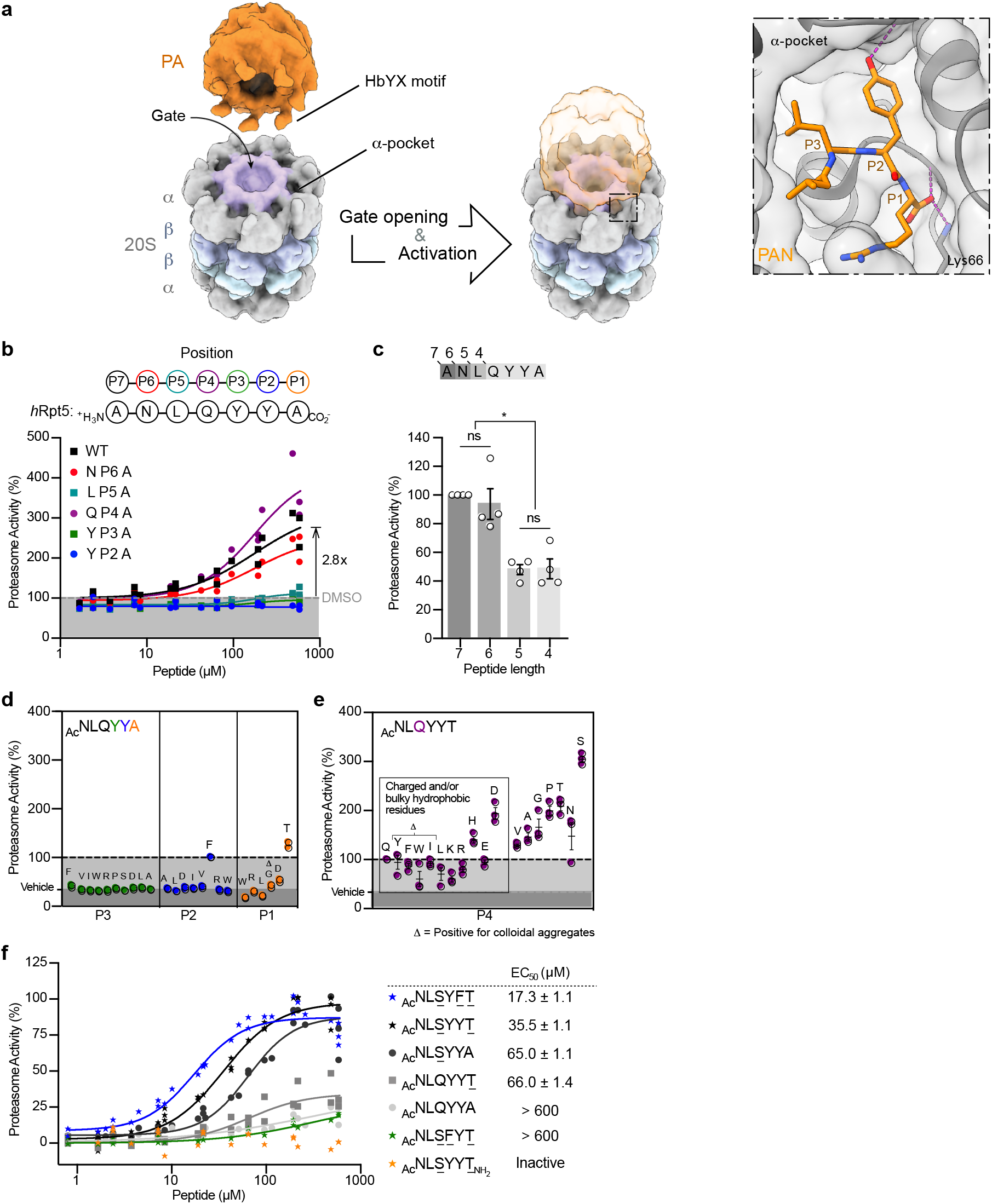
Activation of *h*20S by *h*Rpt5-derived peptides reveals unexpected SAR. **a**, A schematic model of proteasome activation by the C-terminal HbYX motif of a model PA. HbYX motifs are known to dock into α-pockets located between subunits of the 20S α-rings (gray). This interaction opens the gates of the 20S, allowing substrate entry. The inset summarizes the interactions between PAN’s HbYX motif (orange) and the α-pocket (gray surface/cartoon) of the archaeal 20S (PDB ID: 3IPM). **b**, Alanine scanning revealed residues of Rpt5 C-terminus that make key contributions to stimulation of the *h*20S. Activation of the *h*20S (4 nM) was measured by an increase in the hydrolysis rate (RFU/s) of the fluorescent substrate, suc-LLVY-amc (10 µM) upon addition of *h*Rpt5-derived peptides. Data are normalized to DMSO-treated *h*20S and plotted individually (n = 3). **c**, Proteasome stimulation by peptides (250 µM) from an N-terminal truncation series. Results are normalized to ANLQYYA and the average of four independent experiments (open circles; n = 4) is plotted with error reported as s.e.m. P-values were calculated using two-tailed unpaired t-test: *p < 0.04. **d**, Scatterplot of the relative activities of N-terminally acetylated *h*Rpt5 hexapeptides (250 μM) sampling modifications along the P3 (green), P2 (blue) and P1 (orange) residues. Data are normalized to Ac-NLQYYA and plotted individually (n = 2). **e**, Scatterplot of maximal activity of hexapeptides sampling all proteinogenic amino acids at the P4 position (purple). Data are normalized to Ac-NLQYYT and the average of three independent experiments (circles; n = 3) is plotted with error reported as s.e.m. **f**, Combination of optimized residues (underlined) from the SAR campaign. Data are normalized to Opt5 (Ac-NLSYYT, star) and plotted individually (n = 3 or 4). EC_50_ values were calculated from four independent experiments and reported as mean ± s.e.m.

The HbYX model for gate opening was pioneered from studies by Goldberg and colleagues using the C-terminal sequence of the archaeal PA, proteasome-activating nucleotidase (PAN)^9^. This work, and subsequent structures, revealed key roles for the penultimate Tyr residue and the terminal carboxylate of the HbYX motif^12,13^. However, the human 20S proteasome (*h*20S) is a hetero-oligomer instead of the archaeal homo-oligomer, so each of the α-pockets in the *h*20S are distinct. Thus, it is not clear whether the same structure-activity relationships (SAR) that govern recognition of the HbYX motif in the archaeal system are conserved in humans. Addressing this question in the context of natural PAs has been challenging. For example, PA700/19S, the eukaryotic homolog to PAN, limits each of its six distinct C-termini to a specific α-pocket of the 20S^14^, such that contributions of the individual PPIs are likely to be convoluted by limits on the binding topology and cooperativity, along with the confounding effects from ATP hydrolysis^15–17^ and allosteric effects from additional subunits of PA700^18–20^. We envisioned that one way to circumvent these issues might be to use synthetic peptides instead of native PAs. Indeed, HbYX-containing peptides derived from the C-termini of the human Rpt subunits of PA700, have been shown to bind the α-pockets and stimulate turnover of substrates by the 20S^9,21,22^. Inspired by that approach, we hypothesized that a library of peptides could be used to understand the SAR of the HbYX motif in the *h*20S. Importantly, beyond the important insights into the molecular mechanisms of *h*20S gate opening, such investigations might be expected to inform the design of small molecules that bind the α-pockets^23^.

In this study, we designed and synthesized ∼120 peptides derived from Rpt5’s C-terminus and evaluated their ability to stimulate the peptidase activity of the *h*20S *in vitro*. This analysis revealed sequence preferences that differed from the canonical HbYX motif as derived in archaea. We refer to this re-defined preference as the YΦ motif. To better understand the structural underpinnings of the YΦ motif, we grafted an optimal sequence, NLSYYT, to the C-termini of an inert PA platform and solved a 2.9 Å resolution structure of the PA bound to the *h*20S by cryo-electron microscopy (cryo-EM). Remarkably, the same orientation of NLSYYT was observed in five of the seven α-pockets, suggesting a conserved set of molecular interactions. Analysis of the bound YΦ motif revealed specific inter- and intra-molecular contacts, which were involved in molecular recognition and gate opening. Finally, using a series of chimeras, we found that the valence of the PA displaying the YΦ motif (*e*.*g*. monomer *vs*. heptamer) tuned these sequence preferences, with multivalent PAs able to overcome otherwise non-ideal sequences. Together, these studies reveal mechanisms of termini-dependent gate opening in the *h*20S and establish a consensus sequence for monovalent activators.

## RESULTS

### Nomenclature of the HbYX motif and α-pockets

Throughout this work, we will refer to the carboxy-terminal residue of the HbYX motif is termed P1, the next residue P2, *etc*. In this parlance, the canonical HbYX model is defined as having a hydrophobic residue at P3, a preference for Tyr at P2, and any residue at P1. Another key part of the HbYX motif is that it contains the terminal carboxylate at the P1 position, which forms a critical salt bridge with a conserved cationic side chain at the base of the α-pocket; for example αLys66 of the *Thermoplasma acidophilum* 20S (**Fig. 1a**)^9^. Unless otherwise noted, we will use the residue numbering of the *T. acidophilum* 20S.

### Optimization of assay conditions for measuring activation of the h20S

To determine the SAR of the HbYX motif in the *h*20S system, we synthesized, characterized, and assayed a library of peptides derived from the C-terminus of the human Rpt5 (*h*Rpt5), with the native sequence: ANLQYYA. For each peptide, its capacity to accelerate substrate turnover by purified *h*20S was monitored by the hydrolysis of a fluorogenic substrate (suc-LLVY-amc). Because sodium dodecyl sulfate (SDS) and other detergent-like molecules have been reported to non-specifically activate the 20S, these otherwise useful additives are typically excluded from buffers in proteasome activity assays^24^. This constraint makes it more difficult to study activation by peptides, especially if they are minimally soluble or prone to aggregation. Indeed, peptide-based activators have been shown to exhibit atypical dose-responses, including partial to full inhibition of the 20S at higher concentrations^25^.

To minimize these effects, we first optimized the buffer conditions and establish a triage pipeline to remove non-specific activators. These efforts were inspired by previous reports to identify buffers in which the difference between the 20S activities in the basal and stimulated conditions are maximized^5,26,27^. Using this starting condition (see Methods), we observed that high concentrations of either peptide or known small-molecule activators would inhibit, rather than stimulate, substrate turnover by the *h*20S. We reasoned that this effect was likely due to aggregation at the higher concentrations^28^. To solve this issue, we supplemented the assay buffer with a non-ionic surfactant (0.01% Pluronic F-68 ®), which restored a normal dose-responsive curve for most activators without aberrant activation by the detergent itself (**Supplementary Fig. 1a**-**d**). Even under these conditions, we noted that peptides containing thiols (*i*.*e*. Cys and Met) lost their activity over time (**Supplementary Fig. 1e**), even in the presence of dithiothreitol (DTT), so peptides with these residues were excluded from further study. We also omitted peptides that yielded visibly cloudy solutions at 10 mM in DMSO (**Supplementary Fig. 1f**). Finally, the remaining peptides were also subject to analysis by dynamic light scattering (DLS) to detect colloidal aggregation, and any peptides with solubility limits < 600 µM were flagged. Together, these triage steps restricted the sequence space that could be studied, but also minimized contributions from non-specific mechanisms.

### Structure-activity relationships (SAR) of *h*Rpt5 peptides deviate from the HbYX model

In the first series of peptides, we performed an alanine mutational scan of *h*Rpt5: ANLQYYA. Consistent with previous reports^21^, the parent peptide (termed wildtype or WT) stimulated turnover of suc-LLVY-amc by ∼3× (**Fig. 1b**). An EC_50_ value could not be determined, due to relatively weak potency and limited solubility, so, at this stage, we compared peptides based on their relative ability to stimulate hydrolysis rate. Replacing either the P3 (‘Hb’) or P2 (‘Y’) positions with Ala completely ablated activity, in agreement with the HbYX model^9^. Surprisingly, replacing the P4, P5 or P6 residues with Ala also had appreciable effects, including an unexpected enhancement in activity by Ala substitution at P4 (1.3× over WT), complete ablation of activity at P5, and modest impairment at P6 (∼80% of WT) (**Fig. 1b**). A series of N-terminal truncations revealed that the 6-and 7-mer peptides were significantly more stimulatory than shorter ones (**Fig. 1c**), supporting a role for residues “upstream” of the tripeptide motif.^9^

Next, we designed a series of hexapeptides to probe the SAR in more detail. Peptides were synthesized and N-terminally acetylated (Ac), which generally enhanced activity over the free amine (**Supplementary Fig. 2a**). Acetylated peptides were then assessed at a single dose (250 µM) against the *h*20S to obtain an initial overview of their relative activities. In the first set of comparisons, we varied the P1, P2, and P3 positions, replacing the wild-type residue with chemically and/or structurally diverse amino acids. We found that the *h*20S had sequence preferences at the P1 position, with Thr favored over other residues that were tested. More importantly, Tyr seemed to be exclusively required at the P3 position; such that even other hydrophobic residues, including Leu and Ile, were not tolerated. Lastly, either Phe or Tyr was preferred at the P2 position (**Fig. 1d**). When we compared these sequence preferences to those reported for the archaeal 20S, we found that the *h*20S had had both similarities and significant differences (**Supplementary Fig. 3**)^9^.

**Figure 2.**
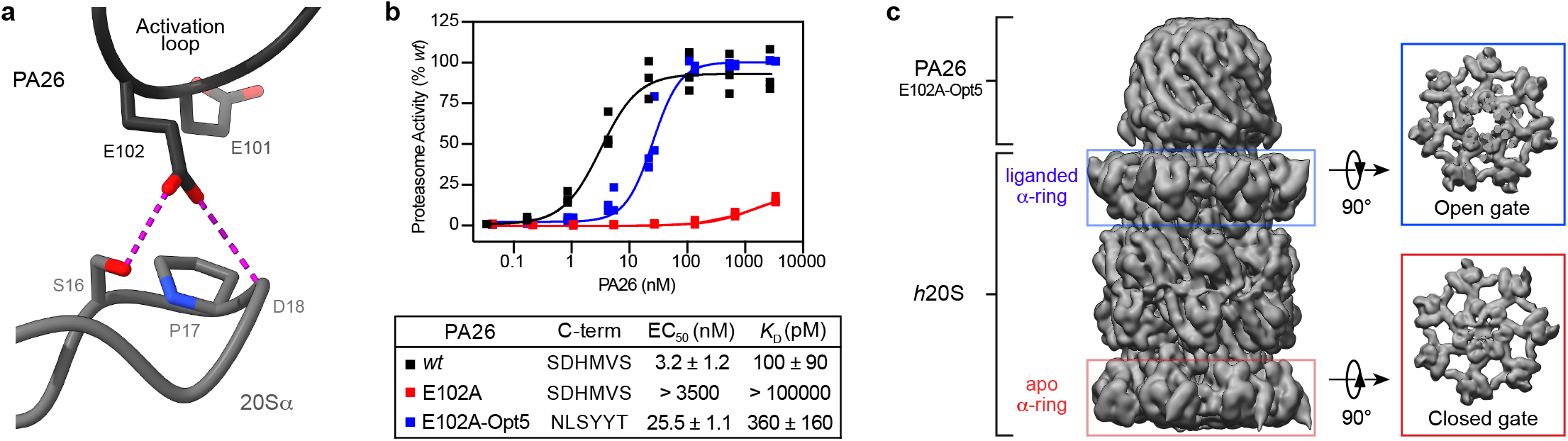
PA26^E102A-Opt5^ induces terminus-dependent activation and gate opening of the *h*20S. **a**, Schematic of how PA26’s activation loop (black cartoon) is known to reposition the Pro17 reverse turn in the 20S α-subunit (gray cartoon) to induce HbYX-independent gate opening (PDB ID: 1YA7). **b**, Stimulation of *h*20S by PA26 constructs: wild type PA26 (*wt*, black), disabled-loop mutant (PA26^E102A^, red), and a disabled-loop mutant with C-terminal Opt5 (NLSYYT) sequence (PA26^E102A-Opt5^), blue). Data are normalized to PA26^E102A-Opt5^ and plotted individually (n = 3). Reported EC_50_ is a mean of EC_50_ values calculated from three independent experiments (n = 3) with error reported as s.e.m. *K*_D_ values were determined by BLI from three independent experiments (n = 3) and reported as the mean ± s.e.m. **c**, PA26^E102A-Opt5^ induces gate opening. The 3D reconstruction of the singly capped PA26^E102A-Opt5^-*h*20S complex at 2.9 Å resolution from 234,960 particles shows gate opening (blue box) relative to the apo, closed-gate α-ring on the opposite side (red box).

**Figure 3.**
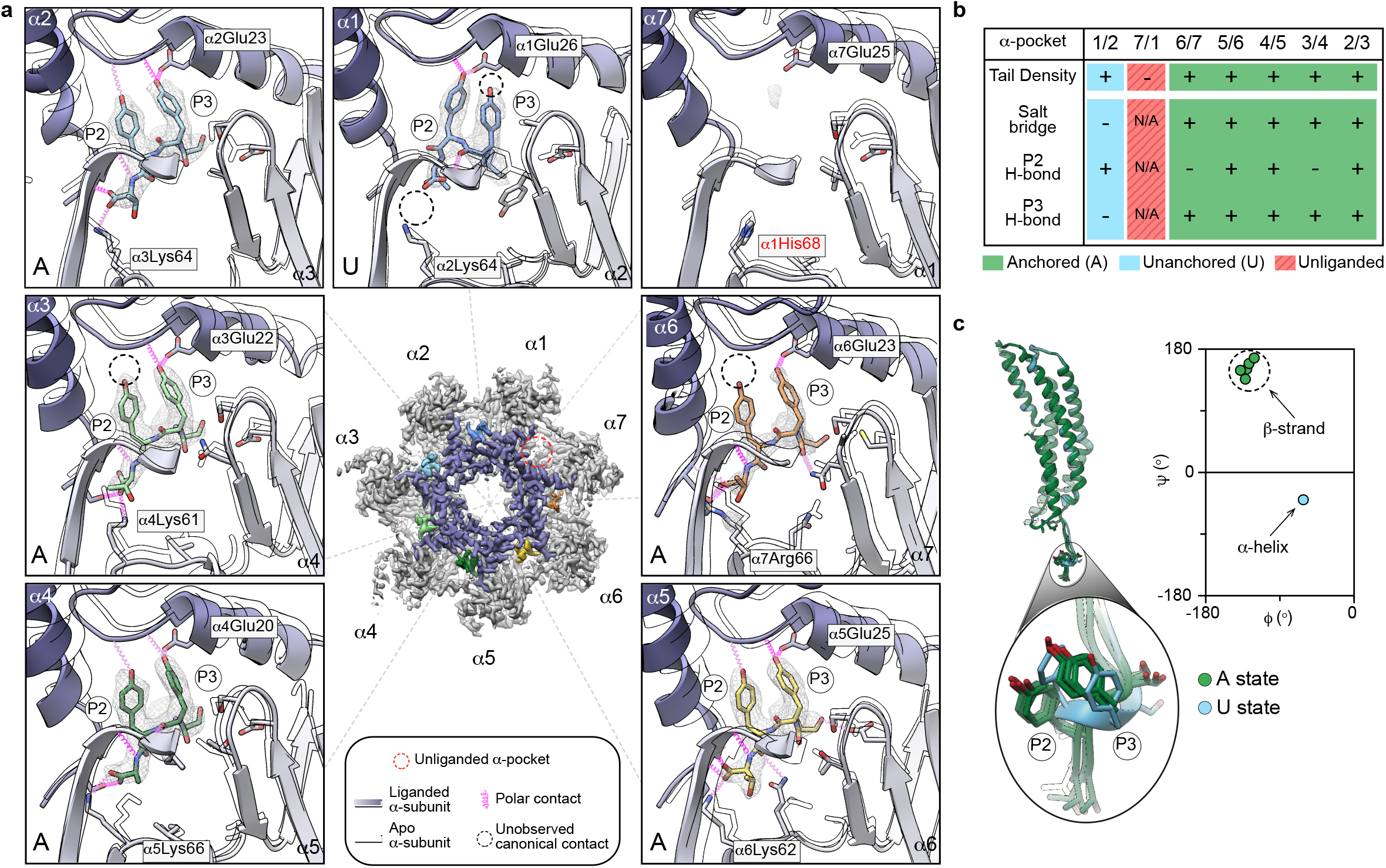
Ensemble of C-terminal interactions promote gate opening in the *h*20S. **a**, Opt5 sequence in the *h*20S α-pockets. The α-subunits are numbered according to the yeast proteasome. *Center*, a top-view 3D density map of the liganded α-ring displaying color-coded densities for each C-tail. The empty α-pocket (α 7/α 1) is marked by a red dotted circle. Close-up views of each C-terminal Opt5 sequence (stick) is superimposed over their corresponding cryo-EM densities (mesh). PA26^E102A-Opt5^ induces terminus-dependent conformational changes throughout the inner (purple) and outer (gray) regions of the α-pockets relative to the apo *h*20S (outlined) (PDB ID: 4R3O). Predicted interactions (pink coils) and unobserved canonical interactions (black dotted circles) are denoted. Anchored (A) or unanchored (U) Opt5 binding states are labeled at the bottom-left corner of each panel (see text). **b**, Summary table of the interactions for each α-pocket. **c**, Opt5 binds with distinct conformations in the A and U states. Opt5 forms an -helix In the U state (α 1/α 2-pocket), while it adopts a β-strand conformation in the A state. Overlaying the C-tails highlights the distinct backbone conformations between the A and U states, while also showing the striking similarities between the A states, especially in the position of the P2 and P3 Tyr sidechains. See **Supplementary Table 2** for relative densities of the C-tails (**b**) and torsion angles (**c**).

We were particularly intrigued by the gain in activity caused by Ala at P4 (see **Fig. 1b**), so we generated a focused series of P4-substituted hexapeptides to explore this position further. In this collection, the P1 was uniformly replaced with Thr, which we found to improve aqueous solubility and activate better than Ala, allowing us to obtain saturable stimulation curves and calculate EC_50_ values (**Supplementary Fig. 4a**). We found that none of the P4 substitutions completely ablated activity, suggesting that, unlike P2 or P3, the requirements at the P4 residue are more permissive. Nonetheless, modifications at P4 impart the greatest modulatory effect on rate of hydrolysis (up to 3× over WT), with positively charged or bulky side chains generally exhibiting lesser stimulation (**Fig. 1e**). Additionally, residues known to disrupt α-helical character (*e*.*g*. Gly and Pro) were well-tolerated at the P4 position, consistent with previous observations^29^. Lastly, we explored the contributions of the P5 and P6 position. Even though we had previously found that truncations of these residues diminished activity, residues substitutions at P5 or P6 had modest effects (see **Supplementary Fig. 2b-d**), suggesting that the side chain identity was less important in these positions.

**Figure 4.**
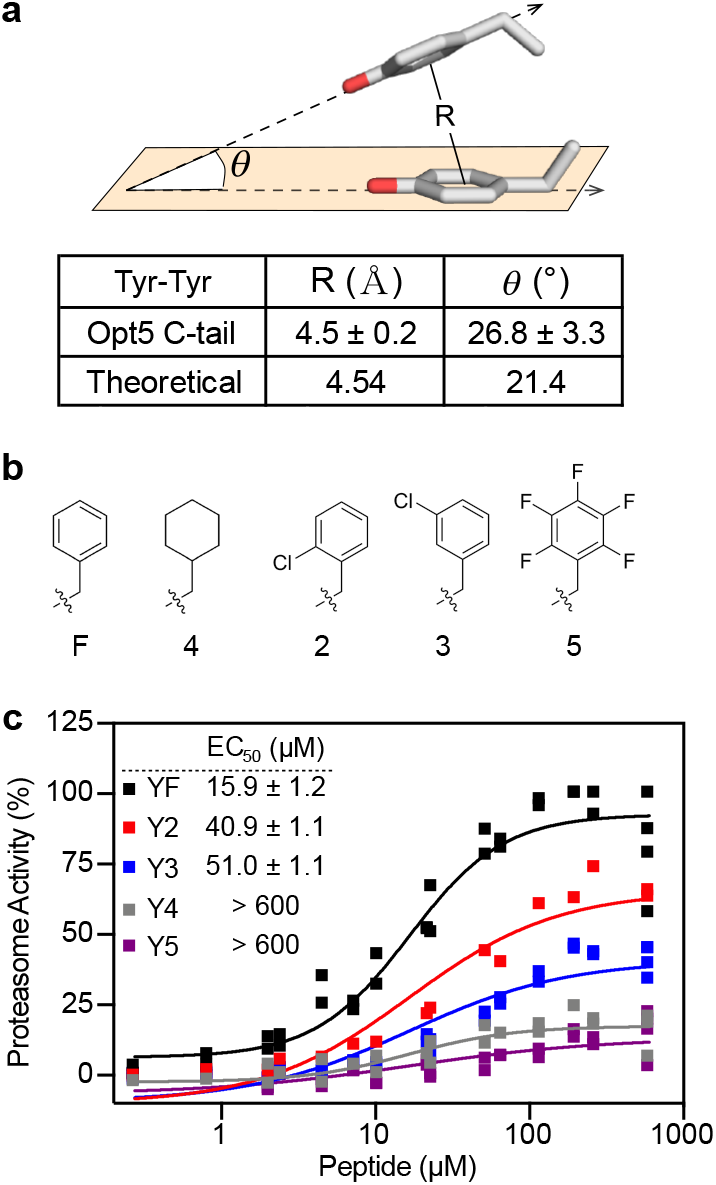
P3 and P2 residues make intramolecular π-stacking interactions within the *h*20S α- pockets. **a** Experimental and theoretical distances (*R*) and angles of incidence (*θ)* for adjacent P2 and P3 tyrosine residues ^38^. Values are of bound Opt5 from the cryo-EM structure (n = 6) and reported as mean ± s.d. See **Supplementary Table 3** for individual measurements. **b** Structure of the side chains from unnatural amino acids used in this study. **c**, P2 modifications to the π electron density decreases the stimulatory effect of Opt5. Data are normalized to Opt5^YF^ and plotted individually (n = 2 to 4). Reported EC_50_ is a mean of EC_50_ values calculated from two to four independent experiments with error reported as s.e.m.

Combining the highlights of the SAR, we incorporated the optimal substitutions at both P1 and P4 to probe whether they are synergistic. In these experiments, we were particularly interested in understanding which substitutions might enhance the apparent EC50 (as a pseudo approximation of affinity) and which ones might impact the rate of hydrolysis. The results showed that Thr at P1 improved only the EC_50_, while Ser at P4 enhanced both EC_50_ and rate of hydrolysis. Combining both substitutions had an additive effect (**Fig. 1f**), yielding an optimized sequence Ac-NLSYYT (EC_50_ = 35.5 ± 1.1 μM). For brevity, we refer to this sequence as Opt5.

Because it seemed potentially distinct from the predictions of the HbYX model, we were interested in the observation that the P2 position accepted either Phe or Tyr (F/Y – denoted as Φ), while Tyr seemed to be critical at the P3 position. To test this idea, we substituted either the P2 or P3 Tyr residues of Opt5 (referred to here as Opt5^YY^ for clarity) with Phe (Opt5^YF^ or Opt5^FY^, respectively) and measured the ability of these peptides to stimulate *h*20S. Removal of the P3 hydroxyl (Opt5^FY^) dramatically reduced activity by ∼70%, while loss of the P2 hydroxyl (Opt5^YF^) had no appreciable effect on turnover and actually improved potency by ∼2-fold relative to Opt5^YY^ (EC_50_ = 17.3 ± 1.1 μM) (**Fig. 1f**). Previous work had also suggested that a Phe at P2 might be favored^30^, so we assessed the generality of this finding by testing two additional peptide sequences (Ac-NLSYΦA and Ac-NLGYΦT), noting up to a 3-fold improvement in potency by Phe over Tyr in both cases (**Supplementary Fig. 4b**).

To this point, measurements of proteasome activity were restricted to the suc-LLVY-amc probe, which measures chymotryptic-like activity. If the optimized peptide Opt5 induces gate opening, we would expect that it would also stimulate the tryptic-like activity. Indeed, we found that treatment with Opt5 and other peptides promoted the tryptic-like activity of the *h*20S, as measured using boc-LRR-amc (**Supplementary Fig. 5a**). Moreover, these peptides also stimulated hydrolysis of a longer, nonapeptide substrate (FAM-LFP), which is known to require gate opening for its entry into the proteasome^31^ (**Supplementary Fig. 5b**). Taken together, these findings suggest that *h*20S activation by *h*Rpt5-like peptides occurs through interactions with at least the last four residues, with additional contributions from P5 and P6. The sequence requirements are expanded from the HbYX motif and are summarized as the YΦ model (see below).

**Figure 5.**
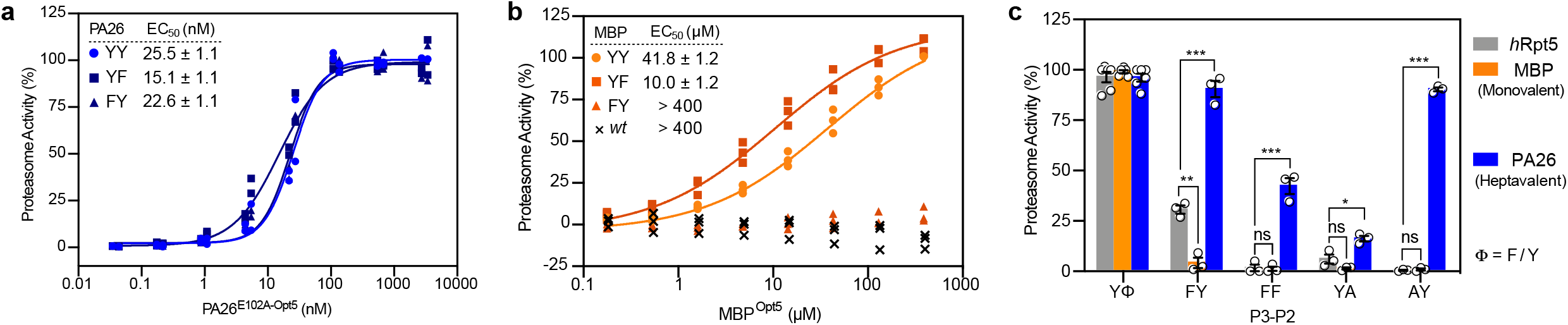
Valency tunes the sequence preferences of *h*20S PAs. **a**, Proteasome activity of heptameric PA26^E102A-Opt5^ (YY, circle) and C-terminally modified variants, YP2F mutant (YF, square) and YP3F mutant (FY, triangle). **b**, Similar to **a** except using C-terminally modified mutants of MBP, tested alongside wildtype MBP (*wt*, ‘x’). Data are normalized to YY and plotted individually (n = 3 or 4). Reported EC_50_ is a mean of EC_50_ values calculated from three independent experiments (n = 3 or 4) with error reported as s.e.m. **c**, Proteasome stimulation by monovalent Opt5 peptides (gray), monovalent MBP^Opt5^ chimeras (orange) and multivalent PA26^E102A-Opt5^ (blue). YΦ is a composite of YY and YF activators. Data are calculated from three independent experiments (see **Supplementary Fig. 4g-h**), normalized to YY, and plotted individually (open circles, n = 6 or 3) with error reported as s.e.m. P-values were calculated using unpaired t-test: *p < 0.05, **p < 0.01, and ***p < 0.001.

### PA26^E102A-Opt5^ induces terminus-dependent gate opening of the *h*20S

To understand the molecular basis for the sequence preferences in the YΦ motif, we attempted to determine the structure of *h*20S bound to Opt5 peptides by cryo-EM. Unfortunately, we observed that these peptides bound with inconsistent occupancy across the α-pockets and we were unable to obtain high resolution structures. Rather, inspired by previous studies^8,13,30,32^, we sought to use the homo-heptameric activator PA26 as a scaffold for the display of multiple copies of Opt5 (NLSYYT). Instead of C-terminal HbYX motifs, PA26 has an activation loop that directly repositions the reverse-turn loop at αPro17 in the α-ring, displacing the N-terminal gating residues (**Fig. 2a**)^3,4^. An Ala substitution in the activation loop (E102A) renders PA26 inactive^8^. Previous studies showed that the capacity of the disabled-loop mutant (PA26^E102A^) to induce gate opening was rescued by grafting PAN’s HbYX motif in place of the native C-termini^13,30^. Thus, we envisioned likewise using PA26^E102A^ as a scaffold to enable high-resolution structural studies.

PA26 from archaea is capable of activating the *h*20S^33^, however, it remains unclear whether the E102A mutation impairs this function, as has been reported in archaea. Using the *T. brucei* PA26, we verified that PA26^E102A^ was unable to stimulate turnover by the *h*20S. Moreover, PA26^E102A^ could no longer bind *h*20S, as measured by biolayer interferometry (BLI) (**Fig. 2b**). Next, we installed Opt5 in place of the last six native residues of PA26 (PA26^E102A-Opt5^) to generate a PA that now bound the *h*20S (*K*_D_ = 360 ± 160 pM) and potently stimulated its activity (EC_50_ = 25.5 ± 1.1 nM) (**Fig. 2b**). To probe how Opt5 binds and activates *h*20S, we then determined the cryo-EM structure of the singly capped *h*20S-PA26^E102A-Opt5^ complex (2.9 Å in overall resolution) (**Supplementary Figs. 6** & **7** and **Supplementary Table 2**). A single state was resolved with the α-rings adjacent to the PA26^E102A-Opt5^ displaying N-terminal extensions that were displaced from the central pore **(Fig. 2c**). At this site, the diameter of the pore was widened by 3.8 Å (**Supplementary Table 3**), reminiscent of other open-gate structures^8^. On the side opposite of the bound PA26^E102A-Opt5^, the gates were relatively closed, consistent with a previous study on the analogous mutant PA26 in complex with the archaeal 20S^32^.

**Figure 6.**
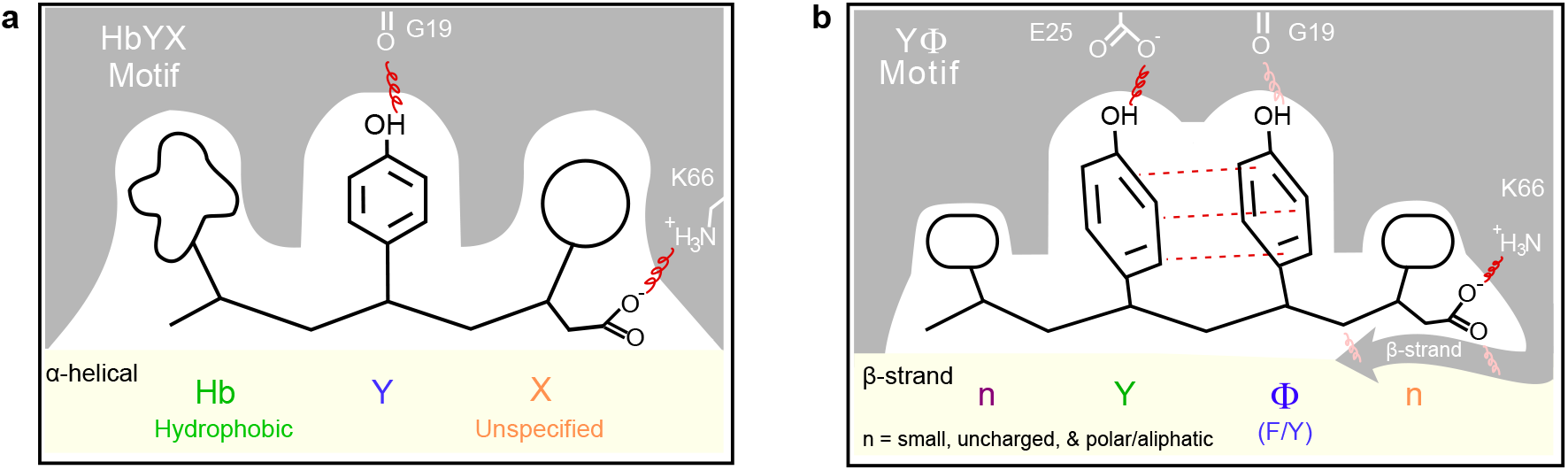
The Y**Φ** motif model for terminus-dependent *h*20S activation. **a**, Model summarizing the major features of the HbYX motif, for comparison. **b**, Model of the nYΦn (YΦ) motif summarizes the proposed mechanism for termini-mediated activation of the *h*20S. Interactions denoted for the Y motif seem specific to monovalent activators of the *h*20S. The critical interactions (red) include polar contacts (squiggle) between Tyr residues and α Glu25 and α Gly19 and pi-stacking between the Tyr residues (dotted line). The carboxylate interaction with α Lys66 is also important. Secondary interactions (pink) are denoted for the P1 (orange), P2 (blue), P3 (green) and P4 (purple) residues. In addition, the P5 and P6 are likely to play a role. Note that multivalency seems to allow PAs to, in some cases, overcome a subset of these requirements.

### Cryo-EM resolves open-gate conformation of the *h*20S

Using this structure, we explored the potential mechanisms by which Opt5 might open the *h*20S gates. In other systems, the open-gate conformation is known to be regulated by a cluster of conserved N-terminal residues (αTyr8, αAsp9, αPro17, and αTyr26)^7,8^. Specifically, these residues are re-positioned away from the central pore during gate opening and are anchored in that position by a characteristic set of intra- and intermolecular contacts. However, these residues are less conserved in humans, particularly in α1 and α2 (α-subunits are labeled to match the numbering used in yeast 20S). We wondered whether these differences might impact the gating mechanism or extent of opening. In the liganded α-ring of our structure, the N-termini of α5, α6, and α7 formed the expected, ordered clusters at the α-subunit interface, consistent with the fully open state^20,34^ and the canonical model. This movement includes the positioning of αPro17 within CH-π distance from αTyr26 and αTyr8. However, the equivalent clusters at the α1/α2 interface were relatively poorly resolved, suggesting a more disordered state, potentially caused by the lack of contacts between the non-canonical α1 Phe8 and α2 Ser9 residues (**Supplementary Fig. 8a**). This “destabilizing effect” also seemed to propagate from the α1/α2 cluster to adjacent α-subunits, resulting in the N-terminal extensions of the α2, α3, and α4 being in a position that partially obstructed the pore, in a conformation that has previously been observed in 26S proteasome structures^16,17^. In those structures, it was not entirely clear whether the C-termini or secondary contacts were responsible for the “partially open” gate conformation, but the observations from this PA26^E102A-Opt5^ complex suggest that termini-mediated mechanisms are sufficient to explain it (**Supplementary Fig. 8b**).

Because the β-rings are known to be relatively unchanged during activation^4,13^, we aligned our structure to these subunits to assess the conformational changes associated with gate opening (PDB ID: 4R3O)^35^. We observed that the Cα atoms of all seven αPro17 residues were radially displaced between 1.0 and 3.7 Å during gate opening. Consistent with the analyses of the N-terminal gate positions, the “partially open” α1- and α2-subunits had the smallest shifts in Pro17 (1.0 and 1.3 Å, respectively) (**Supplementary Fig. 8c**). The modest movement of α2 contrasts to the relatively large shifts typically observed by the α2’s N-terminus in yeast^30^. This is interesting because the yeast α2, like the human α2, also has a non-canonical Ser9; thus, it is possible that α1 Phe8 in the *h*20S, not α2 Ser9, could be more restrictive of the fully open conformation.

We next examined how the termini might be inducing gate opening. First, we confirmed that the disabled activation loop of PA26^E102A-Opt5^ does not interact with αPro17, suggesting that, as intended, this structure allows determination of termini-specific mechanisms. At the α-pockets, most of the P2 and P3 Tyr side chains of bound Opt5 made H-bond contacts with residues that are located in the reverse turn between αPro17 and helix 0 (**Supplementary Fig. 9a**). This finding suggested that residues within Opt5 worked directly through αPro17 to mediate gate opening. As further support of this idea, the α-subunits in these Opt5-bound pockets were rotated an average of 1.5 ± 0.4° about the axial channel (**Supplementary Fig. 9b**). Indeed, previous studies have attributed such rigid body rotation in the α-ring to binding of a C-terminal HbYX motif^12,13,17,32^. In addition, we observed side chain rearrangements throughout the α-pockets that suggest induced-fit conformational changes upon binding of Opt5 (**Fig. 3a**), which has also been implicated in termini-induced gate opening^13^. Overall, we conclude that termini-mediated activation of the *h*20S largely proceeds through the conserved gating mechanism observed in other proteasomes; but with distinctions in the asymmetric gate opening of the α-rings, potentially linked to the non-canonical α1 Phe8.

### Opt5-bound α-pockets reveal key contacts for terminus-dependent *h*20S activation

We next examined the direct interactions of Opt5 in the *h*20S α-pockets. This analysis was aided by the matched stoichiometry of C-termini to α-pockets, together with the fact that the α-pockets were locally resolved to better than 2.9 Å resolution (see **Supplementary Fig. 7d**). Six of the seven α-pockets contained well-defined density corresponding to the termini of PA26^E102A-Opt5^. As previously predicted^8,20,36^, the α7/α1-pocket (interfacing the α7 and α1 subunits) did not contain density for Opt5, likely because this pocket does not have a canonical Lys residue (α1 His68). Of the remaining six sites, the termini in the α1/α2-pocket had a noticeably distinct structure. Specifically, its terminal carboxylate was displaced (5.1 Å) from the ε-amino group of α1 Lys63 and therefore unable to form the expected salt bridge, which has previously been shown to be critical for binding of native PAs and HbYX peptides^9,12^. To verify the importance of the negatively charged terminal carboxylate to overall *h*20S stimulation, we amidated the C-terminus of Opt5 peptide and confirmed that it lost all activity (see **Fig. 1f**). Hence, we categorized the C-termini in the α2/α3, α3/α4, α4/α5, α5/α6 and α6/α7 pockets as binding in an “anchored” (A) state and the C-termini in the α1/α2-pocket as binding in the “unanchored” (U) state (**Fig. 3a, b**).

**Figure 7.**
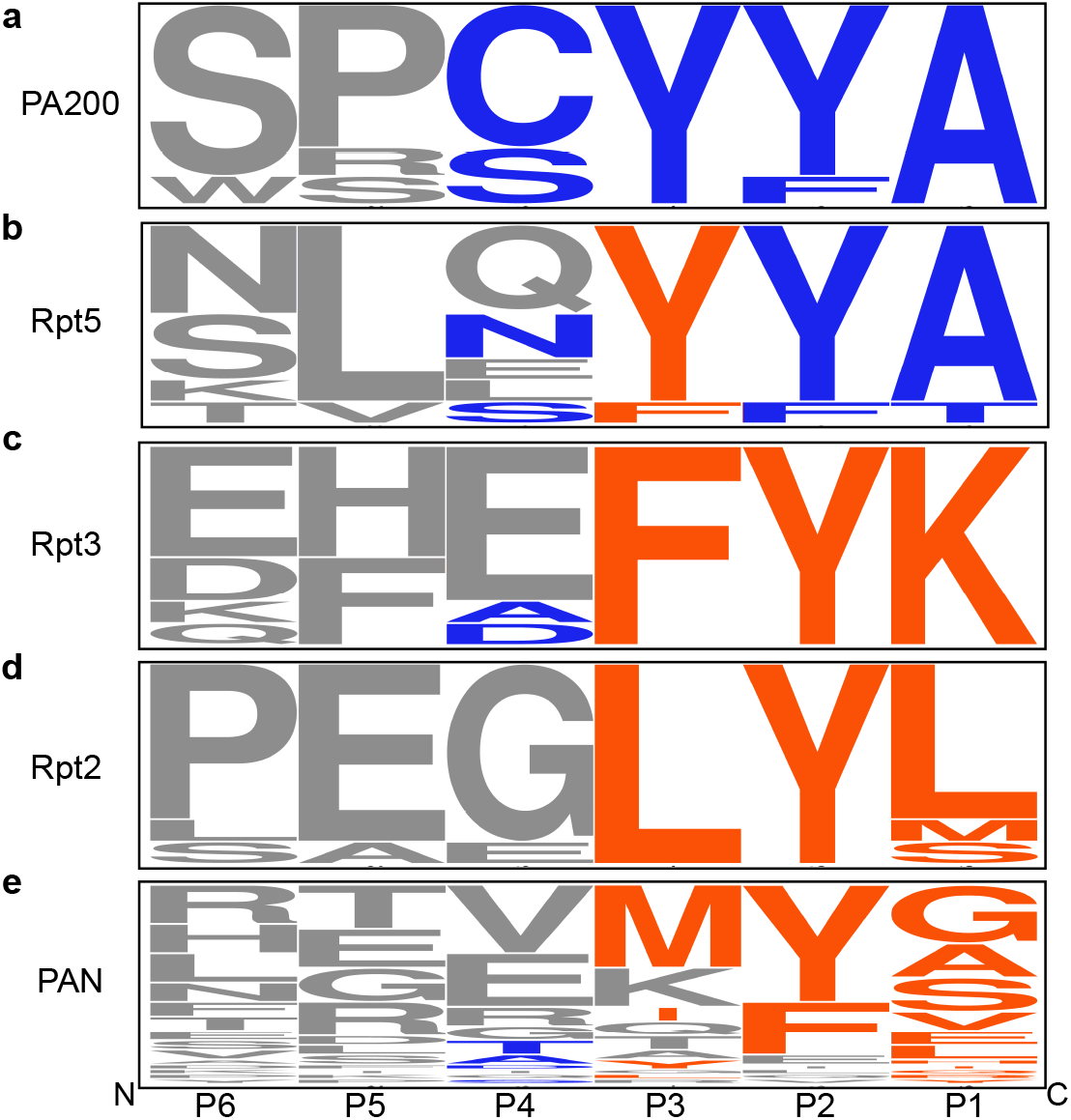
The YΦ motif is strictly conserved in the eukaryotic, monovalent activator PA200. **a-e**, WebLogo (http://weblogo.berkeley.edu) (48) representation of the last six residues of (**a**) PA200 (n = 7), (**b**) Rpt5 (n = 9), (**c**) Rpt3 (n = 9), (**d**) Rpt2 (n = 9) and (**e**) PAN (n = 41). Amino acid residues at the P1, P2, P3 and P4 (Y only) positions are colored according to sequence preferences of the HbYX (orange) or nYΦn (blue) motifs. Unspecified, along with P5 and P6 residues are in gray. The height of each residue represents the relative frequency. The complete entry list in **Supplementary Table 4**.

The peptide backbone of Opt5, of both A and U states, were well-resolved for the P1, P2, P3 and P4 positions, enabling further assessment. First, we noticed that the backbone conformations of the bound C-termini are different between the A and U states. The dihedral angles of the P2-P3 residues of the bound Opt5 form a β-strand structure for the A state but a right-handed α-helix for the U state (**Fig. 3c**). In a recent study of the archaeal 20S-PAN complex, the terminal residues of PAN in PA26^E102A-PAN^ adopt α-helices when docked into α-pockets of the *T. acidophilum* 20S^32^, suggesting that the β-strand conformation might be specific to eukaryotic 20S proteasomes and/or YΦ motifs. Next, we noticed that the five C-termini that bound in the A state are nearly identical; each predicted to make between five and nine hydrogen bonds in the α- pocket, including the critical salt bridge (**Fig. 3a**). In contrast, there are fewer contacts (3 H-bonds and no salt bridge) made between the U state peptide and the α1/α2-pocket. Another distinction between the U and A states is the configuration of predicted H-bonds between Opt5 and residues in the α-pocket, most notably, αGlu25. In the U state, H-bonding occurs between the carboxy side chain of α1 Glu26 and the hydroxyl of the P2 Tyr. In the A state, hydrogen bonding is mediated between αGlu25 and the P3 Tyr side chain, which allows for additional contacts to the α-pocket by the hydroxyl group of the P2 Tyr (*i*.*e*. α2/α3, α4/α5, & α5/α6) (**Fig. 3b**). The αGlu25 is highly conserved from *T. acidophilum* to humans, including across all seven distinct α-subunits (see **Supplementary Fig. 8a**), further highlighting the potential important of this H-bond contact. Together, these studies suggest that polar contacts within the α-pocket, notably H-bonding between the P3 Tyr and αGlu25, are involved in termini-mediated gate opening of the *h*20S.

### Intramolecular stacking of P3 and P2 residues regulates α-pocket engagement

Considering the heterogeneity of the α-pockets in the *h*20S, we were struck by how the C-termini adopted strikingly similar orientations. For example, when we overlay the six bound C-termini from the PA26^E102A-Opt5^-*h*20S structure, the P2 and P3 Tyr residues adopt a nearly identical, stacked orientation (see **Fig. 3c**). Moreover, we noted that a subset of prior structures of eukaryotic PA-20S complexes have C-termini that display a similar stacked arrangement at P2 and P3^10,37^ (**Supplementary Fig. 10**). In our structure, the average distance (*R*) between the centroids of the P2 and P3 Tyr rings (4.5 ± 0.2 Å) and the average angle (*θ)* between the ring planes (26.8 ± 3.3°) closely match theoretical predictions for off-center parallel, π stacking (**Fig. 4a**)^38^. Thus, we postulated that intramolecular π stacking might facilitate termini-dependent gate opening, potentially by orienting Opt5 to a competent conformation within the α-pocket. To test this hypothesis, we designed analogs of the Opt5^YF^ peptide in which π stacking was perturbed and measured their ability to stimulate *h*20S activity. In dose-response assays, activity was reduced 80% relative to Opt5^YF^ when the P2 position was replaced with a cyclohexyl group (Opt5^Y4^), which can no longer π stack with the P3 Tyr side chain (**Fig. 4c**). Likewise, an inversion of the aromatic quadrupole through a pentafluoro phenylalanine (5) substitution at the P2 position (Opt5^Y5^) significantly reduced potency (EC_50_ > 600 µM) and activity (by 80%) relative to Opt5^YF^. Less drastic perturbation of the π electron density, through installation of ortho- or meta-chloro groups (2 and 3; **Fig. 4b**), had intermediate consequences (**Fig. 4c**). Although we cannot rule out steric effects, these experiments suggest that intramolecular π-stacking interactions between the Y and Φ residues contribute to Opt5 activity, perhaps by orienting the motif for engagement with αGlu25.

### Valency tunes the sequence preferences for proteasomal activation

Native PAs and other 20S-binding partners are often multivalent, displaying multiple C-termini from a central scaffold. One effect of this valency is that binding avidity is increased, likely to promote cellular assembly of proteasome complexes^18,26,27,39,40^. However, a less appreciated effect of valency in biological recognition is that it also tunes specificity^41^, amplifying some preferences and allowing more variability in others. To determine if valency influences the sequence preferences of C-terminal recognition by the *h*20S, we generated and tested additional chimeras of the heptavalent PA26^E102A^. In these studies, we focused on the P2 and P3 positions because of their aforementioned, key roles. Unlike the monovalent peptides, Phe substitution of the P3 Tyr did not affect the rate of hydrolysis nor EC_50_ of PA26^E102A-Opt5^ (PA26^YY^ & PA26^FY^; EC_50_ = 25.5 ± 1.1 & 22.6 ± 1.1 nM, respectively). Interestingly, Phe substitution of the P2 Tyr slightly improved the EC_50_ (PA26^YF^; 15.1 ± 1.1 nM) (**Fig. 5a**), consistent with what was observed with the Opt5 peptides. We also noted that PA26^YY^ has a *k*_on_ that is ∼3x slower than that of PA26^YF^ and PA26^FY^; and yet, the calculated *K*_D_ for PA26^YY^ falls within error of the others, owing to its significantly slower *k*_off_ (**Supplementary Fig. 11b**). We then generated an alternate, monovalent PA by fusing peptide sequences onto the C-terminus of maltose-binding protein (MBP). Using these chimeric, monovalent PAs, we found that MBP with a C-terminal Opt5 (MBP^YY^) stimulated the *h*20S (EC_50_ = 41.8 ± 1.2 µM) equipotently to Opt5^YY^ peptide. Likewise, the P2 Phe mutant was more potent (MBP^YF^; EC_50_ = 10.0 ± 1.2 µM), mirroring the SAR from the monovalent peptides. The P3 Phe mutant exhibited diminished activity (MBP^FY^; EC_50_ > 400 µM), however, its relative activity was relatively lower than its peptide equivalent **(Fig. 5b**). Together, we conclude that valency appears to allow the *h*20S to be more permissive of missing contacts in the PA’s C-termini; whereas monovalent ones largely adhere to the YΦ motif.

Next, we interrogated the impact valency might have on the relative importance of intramolecular π stacking by generating P3 or P2 Ala mutations in all three PA-types: monovalent peptide, monovalent MBP, and multivalent PA26^E102A^. In stimulation assays, Ala mutations at either P3 or P2 completely inactivated the monomeric tools: peptides (Opt5^AY^ and Opt5^YA^) and MBP chimeras (MBP^AY^ and MBP^YA^) (**Fig 5c**; **Supplementary Fig. 4d, g**). Thus, without intramolecular π stacking, monovalent PAs seem unable to stimulate *h*20S. However, in the context of multivalent PA26^E102A^, disrupting π stacking with a P3 Ala mutation (PA26^AY^) had no effect on hydrolysis (full hydrolysis and EC_50_ ∼ 8.3 nM). Rather, introduction of the P2 Ala mutation (PA26^YA^) was required to ablate stimulation (partial activation and EC_50_ > 3000 nM; **Supplementary Fig. 4d**). Thus, multivalent PA’s do not seem to require either P3 Tyr or intramolecular π stacking to induce gate opening. Another distinction between mono- and multi-valent activators is the residue preferences at the P1 and P4 positions, which were effectively masked in the heptavalent PA26^E102A^. For example, PA26^NLQYYA^ activates more potently (EC_50_ = 6.2 ± 1.1 nM) than the peptide SAR-derived, Opt5 sequence (**Supplementary Fig. 4d**). Collectively, these data show that valency reduces the stringency for termini-mediated activation of the *h*20S (**Fig. 5c**).

## DISCUSSION

### An updated HbYX model for binding the α-pockets and initiating *h*20S gate opening

Pioneering studies of the HbYX motif in the archaeal system (**Fig. 6a**)^9,12,13^, have contributed significantly to our understanding of 20S activation. Herein, we aimed to deepen that knowledge through characterization of how Rpt5-derived peptides stimulate the *h*20S. This effort uncovered narrower preferences for the P1 ‘X’ and P3 ‘hydrophobic’ residues and revealed an unanticipated role for the P4 position, suggesting that the *h*20S exhibits distinct sequence preferences. Although only a subset of amino acids could be explored because of solubility criteria, only Ala and Thr were strongly preferred at P1, a result that contrasts with the relatively broad specificity exhibited in the archaeal system (see **Supplementary Fig. 3**). A potential reason for this preference was revealed by structural studies, which showed that Thr in the P1 position engages in H-bonding within *h*20S α-pockets. In contrast, the structural rationale for the preference at P4 is less clear. Tentatively, small, aliphatic residues might be preferred over bulkier residues due to potential steric clashes with a neighboring β-strand-turn in the α-pockets. In addition, a subset of polar amino acids, such as Ser, Thr, and Asp, were also relatively preferred at P4 (see **Fig. 1e**), indicating that H-bonding in that region may also be involved. However, further work is needed to elucidate the importance of these interactions, including whether they contribute to binding, gate opening or both. Finally, another key observation was that a Tyr was prioritized at P3, whereas the P2 position accepted Φ. Structural and activity analyses suggested that these preferences might be based on a requirement for intermolecular H-bonds with the α-pocket (esp. a conserved glutamate) and intramolecular π-stacking interactions. Together, these structure-function observations led us to propose the YΦ motif as an addendum to the HbYX model, which is focused on the *h*20S (**Fig. 6b**).

In our *h*20S-PA26^E102A-Opt5^ structure, we noted that the C-termini bind in two different states (A and U). Although Opt peptide is monomeric (*e*.*g*. not attached to PA26), we speculate that it might occupy the α-pockets in conformations that resemble A and/or U, such that the structure could be informative of key contacts. Specifically, we speculate that the A and U states potentially represents putative docking poses for Opt5^YΦ^ (Opt5^YF^ and Opt5^YY^) and Opt5^FY^ peptides, respectively. For example, the diminished activity of Opt5^FY^ might be linked to fewer H-bond contacts and the displaced salt bridge depicted in the U state, because the P3 Phe is unable to engage with the αGlu25, such that the P3 Tyr would have to “reach over” to this position (as in the U state). Similarly, Opt5^YΦ^ peptides that bind in an orientation that resembles the A state would be expected to form more contacts in the α-pocket, including the salt bridge and H-bonds between the P3 Tyr and αGlu25. In reported structures of both yeast^20^ and human^37^ 26S proteasomes, the C-termini of the Rpt3 subunit binds the α1/α2-pocket with a displaced salt bridge while adopting an α-helical turn, as observed in the U state. Moreover, the P3-P2 residues of Rpt3’s C-terminal consensus sequence are, like Opt5^FY^, F-Y, which we predicted to be associated with the U state. This speculation aside, it is also possible that the observed U state is entirely independent of Opt5 and its interactions and instead stems from unique features of the α1/α2-pocket. Additional structures, particularly of peptide-bound complexes, will be required to deepen our understanding of the YΦ gating mechanism.

### Valency tunes selectivity for the C-terminal PA sequences

Using a series of monovalent (free peptides and MBP chimeras) and heptameric (PA26^E102A^) scaffolds, we explored how valency tunes sequence preferences. We found that the multivalent PA26 ^E102A^ was able to overcome a subset of the requirements suggested by the YΦ motif, most strikingly, a Tyr residue at the P3 position. This finding implies that monovalent, eukaryotic activators, such as PA200/Blm10^10^, might have distinct requirements at their C-termini. As a first step in asking that question, we aligned the C-terminal sequences of naturally occurring PAs from multiple organisms (**Supplementary Table 4**). Indeed, PA200 ended in residues that closely match the YΦ motif (**Fig. 7a**). Moreover, a recent structure revealed that the C-terminus of PA200 adopts both the β-strand and the intramolecular off-centered π stacking^42^. Thus, we postulate that monovalent PAs will strictly adhere to the YΦ model. A different picture emerged from alignment of C-terminal sequences from multivalent PAs, including archaeal PAN and the three Rpt subunits of PA700 containing HbYX motifs (Rpt2, Rpt3, & Rpt5) from multiple organisms. This exercise showed that the C-terminus of Rpt5 closely resembled the YΦ motif, except that the P3 position accommodates a Phe residue, and the P4 position samples a wider range of residues (**Fig. 7b**). Notably, Rpt5 is the most stimulatory of the Rpt subunits^21,22^, perhaps supporting the influence of the YΦ motif on *h*20S activation, even in the context of some multivalent PAs. The less stimulatory Rpt2 and Rpt3 subunits, in contrast, displayed more divergent sequences from the YΦ motif, most notably because Tyr was not conserved at the P3 position (**Fig. 7c, d**). Even less consensus was observed when we aligned the C-termini of available PAN sequences. Specifically, the P2 tended to be Φ, while the residues at the remaining positions seemed arbitrary (**Fig. 7e**). Unlike PA700, PAN is homo-oligomeric and so no individual monomer (*e*.*g*. Rpt5) is predicted to make an out-sized contribution to overall binding or stimulation. Thus, in the case of PAN, valency seems to largely override the requirements of both the YΦ and the more permissive HbYX motifs. Non-adherence to the canonical HbYX model was similarly noted in an examination of another hexavalent regulator of the archaeal 20S, Cdc48^40^. Together, these observations suggest that C-terminal sequence identity, plus a contribution from valency, combine to dictate the PPIs between PAs and the proteasome. Further analysis of the SAR at these PPIs is important because additional binding partners of the 20S, with HbYX-like motifs of varying sequences and valences, have been identified in eukaryotes, but the full extent to which they regulate the proteasome remains limited^40,43–45^. Hence, it may be useful to create sub-categories of HbYX-like models, in which the valency of the PA is a key, defining characteristic.

### Implications for the discovery of pharmacological proteasome activators

Loss of proteasome function is implicated in many devasting proteinopathies, including neurodegenerative disorders^46^. Consistent with this idea, boosting 20S activity, by introducing either native^47^ or engineered activators^48^, has been found to be protective in cell-based disease models. These observations have motivated campaigns to discover drug-like molecules that mimic the activity of PAs^23^. This current work could be important in that effort, by defining the SAR associated with *h*20S α-pockets and providing a template for the rational design of pharmacological proteasome activators^49,50^. For example, we suspect that arrangement of the YΦ motif, through π stacking, minimizes the entropic costs of binding for Opt5. For molecules that conform to this pharmacophore model, potency should be improved by rigidification of the bioisosteric equivalent to the Tyr-Tyr π stack. We also found that Opt5 docked into 6 of the 7 available α-pockets with remarkable similar structural states. This consensus structure might be a good starting point for understanding the requisite occupancy and binding interactions for inducing gate opening with a small molecule. Thus, the current work provides a potential blueprint for designing novel proteasome activators, while also extending our knowledge of the mechanisms of *h*20S activation.

## SUPPLEMENTARY DATA

**Supplementary Fig. 1. a**, Pipeline for assay optimization and hit validation of peptide activators of the *h*20S. See text for details. Briefly, we started with a buffer previously reported to have optimized dynamic range, and then screened detergents to identify those that would not promote activity, but still minimize artifacts from aggregators. Finally, we measured both the effects on turnover rate (ΔVmax) and the EC_50_ (potency). **b**, Structures of previously reported 20S activators and peptide sequences used as benchmarks in this study. **c**, Addition of surfactant (0.01% Pluronic F-68 ®) restores dose-dependence of *h*20S activation. Representative dose-response curves for small molecule (CPZ, blue; MK-886, green) and *h*Rpt5 peptide (black) activators. Below each graph is a zoom-in is shown to highlight the non-specific inhibition at higher concentrations (above ∼100 µM). We contribute this loss of activity to insolubility. Addition of detergent restores normal activity, except for the small molecule MK-886, which loses its activity (green). **d**, ATP inhibits activation by *h*Rpt5-based peptides. Representative dose-response curve of activation with ATP titration. Based on these results, we selected an assay buffer that contains 200 μM of ATP. **e**, Time-dependent loss of activity in Cys- and Met-containing peptides. Two separate dose-response curves from P4-scan (see **Supplementary Fig. 4b**). **c**-**e**, Technical replicates are plotted individually (n = 2). **f**, Typical, soluble peptide appears clear as 10 mM stock in DMSO (left, Opt5), while aggregated peptide appears white and cloudy (right, *h*Rpt5-P2F).

**Supplementary Fig. 2**. Effects of N-terminal modifications on stimulation of *h*20S by peptides. **a**, N-terminal acetylation of *h*Rpt5-derived peptides tends to modestly enhance their stimulatory activity. The effect of N-terminal acetylation is plotted as log_2_ fold increase (blue) or decrease (red) in proteasomal stimulation. Additional modifications from the wild type *h*Rpt5 sequence are underlined. Reported data is a mean calculated from two independent experiments with error reported as s.e.m. **b-d**, Scatterplots of the relative activities of peptides (250 μM) sampling residues at the P6 (orange) and P5 (teal) positions. Activities are benchmarked by vehicle and either (**b**) Ac-NLQYYA, (**c**) NLGYYA, or (**d**) Ac-NLSYYT (dotted lines). Data are normalized to Ac-NLQYYA and plotted individually (n = 2) along with the mean ± s.e.m.

**Supplementary Fig. 3**. The structure-activity profile for *h*Rpt5-activated *h*20S differs from the HbYX model **a**, Summary of the previously described SAR for PAN-stimulated activity of the Thermoplasma 20S proteasome^9^ (orange). **b**, The SAR for *h*Rpt5-stimulated activity of the human 20S (blue). The corresponding mutations from Smith et al. 2007 were made to Ac-NLQYYA and the data aligned to the archaeal heatmaps. Only a subset of the *h*20S results are included to allow direct comparisons to exact sequences. Data are normalized to WT and plotted as mean (n = 2). **a**,**b**, Residues of wildtype sequences are denoted with an ‘×’. Activity measurements were not determined for residues in gray.

**Supplementary Fig. 4**. Dose-dependent stimulation induced by different PAs. **a-c**, Stimulation of *h*20S by P4-substituted hexapeptides. (**a**) Normalized dose-response curves. Data are normalized to Opt5 (Ac-NLSYYT) and plotted individually (n = 3). (**b**) Experimental replicates of non-normalized dose-response curves with technical replicate plotted individually (n=2). (**c**) EC_50_ values are reported for each independent experiment (n = 3). The mean of the “raw” EC_50_ values (yellow) along with the mean of the normalized EC_50_ values (blue) are reported with error as s.e.m. (n = 3). **d**. Dose-response curves for additional *h*Rpt5-derived hexapeptides. Data are normalized to Opt5 and plotted individually (n = 3) with calculated EC_50_ values with error reported as s.e.m. (n = 3). **e**,**f**, Activity of hexapeptides for pi-stacking studies. (**e**) Experimental replicates of raw dose-response curves with technical replicate plotted individually (n=2). (**f**) EC_50_ values are reported for each independent experiment (n = 4). The mean of the “raw” EC_50_ values (yellow) along with the mean of the normalized EC_50_ values (blue) are reported with error as s.e.m. (n = 2 to 4). **g**,**h**, Dose-response curves for additional (**g**) C-terminal MBP mutants and (**h**) C-terminal PA26^E102A^ mutants. Data are normalized to Opt5 sequence and plotted individually (n = 3). Reported EC_50_ is a mean of EC_50_ values calculated from two to four independent experiments with error reported as s.e.m.

**Supplementary Fig. 5**. Distinct proteasome substrates report similar SAR for *h*Rpt5-based activators. **a**,**b**, Stimulation of the *h*20S (4 nM) by *h*Rpt5-based peptides was assessed with (**a**) the trypsin-targeted boc-LRR-amc (20 μM) and (**b**) the gate-sensitive nonapeptide FAM-LFP (100 nM) as alternate substrates to suc-LLVY-amc. Data are normalized to (**a**) Ac-NLSYFT or (**b**) Ac-NLSYYT and plotted individually (n = 2). **c, d** EC_50_ values determined with LRR and LFP substrates were calculated from two independent experiments and reported as the mean ± s.e.m. Corresponding values for LLVY were also included. **d**-**f**, Experimental replicates of non-normalized dose-response curves for optimization panel of *h*Rpt5 assessing hydrolysis of (**d**) LLVY, (**e**) LRR, and (**f**) LFP substrates. Technical replicates (n=2) are plotted individually for two to four independent experiments with the EC_50_ values ± s.e.m. reported for each replicate. The “raw” mean (yellow) EC_50_ was calculated and reported alongside the “normalized” mean (blue) with error reported as s.e.m. (n = 2 or 4).

**Supplementary Fig. 6**. Schematic for cryo-EM single-particle data processing. Low resolution or artefactual reconstructions (white) and corresponding particles were excluded from subsequent processing steps, whereas all other reconstructions (gray) and corresponding particles were utilized. Representative 2D classes (dark circular background) are shown.

**Supplementary Fig. 7**. Cryo-EM metrics for PA26^E102A-Opt5^-*h*20S complex. **a**, Representative micrograph. **b**, The Fourier Shell Correlation (FSC) curves for the unmasked (green), masked (blue) and phase randomized (red) reconstructions. The resolution at 0.143 is indicated by a dashed line. **c**, Distribution of Euler angles displaying the orthogonal view of the reconstruction. The angular distribution is shown as a column whose longitudinal axis aligns with the normal of the corresponding back-projection **d**, Local resolution estimates of the global reconstruction for the complete volume (left) and a coronal cross-section (right). **c**,**d** Generated and calculated with RELION^52,53^.

**Supplementary Fig. 8**. Asymmetric *h*20S open-gate conformation. **a**, N-terminal sequence alignment of 20S proteasome α-subunits from *H. sapien* (Hs) and *T. acidophilum* (Ta). The α-subunits are labeled with human gene name but numbered according to yeast α-subunit/yeast gene. The α-subunits that directly gate the α-annulus (red) with their relative contributions to gating denoted with a gradient. The most N-terminal residue with resolvable backbone density in the cryo-EM reconstruction is underlined. The conserved cluster (yellow), non-canonical residues (gray), and conserved α-pocket residues (purple) are highlighted. **b**, Conformational state of N-terminal gate (yellow) and interactions of conserved cluster (α1, orange; α2, α4, α6, pink; α3, α5, α7, cyan) that stabilize asymmetric open-gate *h*20S. Non-canonical residues (gray) are denoted for resolved residues. Single or double asterisks indicates α-subunits featuring unresolved extreme N-termini with non-canonical (*) or conserved (**) residues, respectively. **c**, Radial displacement of αPro17. Individual measurements are reported along with mean ± s.d.

**Supplementary Fig. 9**. Structural evidence implicates termini-mediated gate opening. **a**, Disabled activation loop of PA26^E102A-Opt5^ no longer interacts with αPro17 reverse turn in representative α-pocket (α5/α6-pocket). C-terminal Opt5 engages residues within reverse-turn loop from the α-pocket. **b**, PA26^E102A-Opt5^ induces rotation of the α-subunits about the axial channel. Individual measurements are reported along with mean ± s.d.

**Supplementary Fig. 10**. Binding orientation of reported YΦ motifs align with Opt5. **a**,**b**, Overlay highlights similar backbone conformation between bound Opt5 (SYYT, grey) and the C-terminus of (**a**) the human Rpt5 (QYYA, orange) (PDB ID: 5GJR) or (**b**) the yeast Blm10/PA200 (SYYA, green) (PDB ID: 4V70). Aromatic P2 and P3 Tyr residues share π-stacking orientation. Interactions with α-pocket residues are denoted. For *h*20S structure, use PDB ID: 6REY…

**Supplementary Fig. 11**. Biolayer interferometry (BLI) of PA26 activators. **a**, Sensorgram traces of *h*20S binding to either wildtype (*wt*, red) or E102A (blue) variants of PA26 at different concentrations. Colored curves represent measurements for individual experiments (n = 2 or 3) and solid black lines correspond to global-fit for a representative experiment. **b**, Binding and kinetic constants measured by BLI for binding of PA26 to *h*20S. N.D. denotes values that could not be determined due to undetectable binding.

**Supplementary Table 1**. *h*Rpt5-derived peptides

**Supplementary Table 2**. Cryo-EM data collection, refinement, and validation statistics

**Supplementary Table 3**. Measurements associated with open-gate *h*20S cryo-EM structure

**Supplementary Table 4**. C-terminal sequences of PA200, PAN, Rpt5, Rpt3 and Rpt2 from different organisms identified in UniProtKB.

## METHODS

### Reagents

Human 20S proteasome was purchased from the Proteasome Center. pET28 6His PA26 (49V) cloning vector was a gift from Philip Coffino. pET 6His MBP TEV LIC cloning vector (1M) was a gift from Scott Gradia (Addgene plasmid # 29656).

### Strains and Plasmids

The *E. coli* strain Top10 was used for propagating plasmids. BL21 (DE3) cells were used for expression and purification of recombinant proteins.

### Peptide synthesis

Peptides were synthesized by Fmoc solid phase peptide synthesis on a Syro II peptide synthesizer (Biotage) at ambient temperature and atmosphere on a 12.5 μM using either pre-loaded Wang resin or Rink amide resin (Sigma-Aldrich). Coupling reactions were run with 4.9 eq. of HCTU (*O*-(1H-6-chlorobenzotriazole-1-yl)-1,1,3,3-tetramethyluronium hexafluoro-phosphate), 5 eq. of Fmoc-AA-OH and 20 eq. of *N*-methylmorpholine (NMM) in 500 μl of *N,N*-dimethyl formamide (DMF). Fmoc-AA-OH was double coupled for 8 min while shaking for each position. Fmoc deprotection was conducted with 500 μl 40% 4-methylpiperidine in DMF for 3 min, followed by 500 μl 20% 4-methylpiperidine in DMF for 10 min and six washes with 500 μl of DMF for 3 min. Acetylation of the N-terminus was achieved by reacting 20 eq. acetic anhydride and 20 eq. *N,N*-diisopropylethylamine (DIPEA) in 1 mL DMF for 2 h while shaking. Peptides were cleaved with 500 μl of cleavage solution (95% trifluoroacetic acid (TFA), 2.5% water and 2.5% triisopropylsilane) while shaking for 2 h. Crude peptides were precipitated in 15 ml cold 1:1 diethyl ether: hexanes and air-dried overnight. Crude peptides were solubilized in a 1:1:1 mixture of DMSO: water: acetonitrile, filtered, and purified by high-performance liquid chromatography (HPLC) on an Agilent Pursuit 5 C18 column (5 mm bead size, 150 × 21.2 mm) using Agilent PrepStar 218 series preparative HPLC. The mobile phase consisted of A, 0.1% TFA in water and B, 0.1% TFA in acetonitrile. Peptides were purified to >95% homogeneity confirmed by liquid chromatography-mass spectrometry before solvent was removed by lyophilization. Peptides were resuspended in 1:1 water: acetonitrile, lyophilized again in tared tubes and stocks were stored at -20 °C. As mentioned in the text and Supplemental Fig 1, insoluble peptides were identified by visual inspection, DLS, changing potency over storage time and/or the appearance of atypical concentration curves. Such peptides were excluded from further analysis.

### Protein expression and purification in bacteria

All proteins were produced in *E. coli* BL21(DE3) and stored at −80 °C.

PA26 WT and PA26^E102A^ (*Trypanosoma brucei*, 49V, His tagged) were expressed from a pET28 construct with a N-terminal 6His tag. *E. coli* were grown in terrific broth (TB) with the requisite antibiotic at 37 °C, induced with 1 mM IPTG in log phase, and grown for an additional 3 h. Cells were harvested by centrifugation, resuspended in His resin binding buffer (20 mM Tris pH 7.9, 20 mM NaCl, 10 mM imidazole) supplemented with protease inhibitors, sonicated and clarified by centrifugation. Clarified lysate was applied to Ni-NTA His-Bind Resin (Novagen). Resin was washed with binding buffer and then eluted in batches with buffer containing increasing amounts of imidazole (up to 300 mM). Purified protein was treated with 1 mM DTT for 1 h at ambient temperature and imidazole was removed by overnight dialysis into storage buffer (20 nM Tris pH 8.0, 200 mM NaCl) for later use.

Chimeric PA26^E102A^ (*Trypanosoma brucei*, 49V, His tagged) were expressed from a pMCSG7 construct with an N-terminal TEV-cleavable 6His tag. *E. coli* were grown in TB with the requisite antibiotic at 37 °C, induced with 1 mM IPTG in log phase, cooled to 18 °C and grown overnight. Cells were harvested by centrifugation, resuspended in His resin binding buffer (20 mM Tris pH 7.9, 20 mM NaCl, 10 mM imidazole) supplemented with protease inhibitors, sonicated and clarified by centrifugation. Clarified lysate was applied to Ni-NTA His-Bind Resin (Novagen). Resin was washed with binding buffer and then eluted in batches with buffer containing increasing amounts of imidazole (up to 800 mM). Purified protein was treated with 1 mM DTT for 1 hr at ambient temperature and imidazole was removed by overnight dialysis into storage buffer for later use.

MBP WT and chimeric MBP (His tagged) were expressed from pET and pMCSG7 constructs, respectively. *E. coli* were grown in TB supplemented with 0.2% glucose (instead of glycerol) with the requisite antibiotic at 37 °C, induced with 1 mM IPTG in log phase, cooled to 25 °C and grown overnight. Cells were harvested by centrifugation, resuspended in amylose resin binding buffer (20 mM Tris pH 7.8, 200 mM NaCl, 1 mM EDTA) supplemented with protease inhibitors, sonicated and clarified by centrifugation. Clarified lysate was applied to amylose resin (New England BioLabs, E8021S). Resin was washed with binding buffer and then eluted with 200 mM maltose in binding buffer. Purified protein was dialyzed into storage buffer for later use.

### Proteasome activity assays

Stimulation of *h*20S proteasome (Proteasome Center) was assayed in 384-well plates (Greiner Bio-One, 781209) using the fluorogenic peptide substrates suc-LLVY-amc (AnaSpec, AS-63892), boc-LRR-amc (AdipoGen, AG-CP3-0014) or FAM-LFP (5-FAM-AKVYPYPMEK(QXL520)-NH2; AnaSpec) in assay buffer containing 50 mM Tris pH 7.5,10 mM MgCl_2_, 200 μM ATP, 1 mM DTT, and 0.01% Pluronic F-68® (Gibco Life Technologies, 24040032) in a total volume of 30 μL. Human 20S (final concentration of 4 nM) was incubated in the presence or absence of activators (peptides, 0.2 – 600 μM; MBP 0.2 – 400 μM; PA26, 0.04 – 3500 nM) at ambient temperature for 5 min. Substrate (suc-LLVY-amc, 10 μM; boc-LRR-amc, 20 μM; or FAM-LFP, 100 nM) was added immediately before reading. Fluorescence intensity of suc-LLVY-amc and boc-LRR-amc (excitation, 355 nm; emission, 440 nm; cutoff 435 nm) or FAM-LFP (excitation, 490 nm; emission, 520 nm; cutoff, 515 nm) were monitored at 30 - 60 s intervals for 30 min at 25 °C using a Spectramax M5 microplate reader (Molecular Devices). The hydrolysis rate was calculated from the slope of the curve between 100 and 500 s (500 and 1000 s for boc-LRR-amc) in arbitrary units (RFU per s).

Data were processed and fit in GraphPad Prism 8.0. The baseline hydrolysis was normalized to the total mean activity for the lowest concentration of every activator assayed in a given plate and the maximal hydrolysis was normalized as reported. Normalized activity were plotted relative to log_10_(activator). Data was fit to the model for log(agonist) versus response (variable slope). In equation (1), *X* = log[activator, μM] except for PA26-based activators, which are in nanomolar.

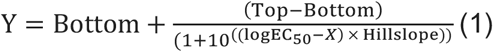

### Dynamic Light Scattering

From 10 mM stocks in DMSO, three 4-fold serial dilutions of each peptide were prepared in assay buffer that was filtered with Millex-GS 0.22 μm sterile filter unit to final concentrations of 600, 150, and 37.5 μM. A final volume of 20 μl was added to a 384-well plate (Corning, 3540) and the unsealed plate was centrifuged at 25 °C for 45 s at 500 x g. Colloidal aggregation was measured using a Wyatt Technologies DynaPro Plate Reader II (acquisition time of 2 s, 10 acquisitions, with auto attenuation, and temperature controlled at 25 °C).

### Biotinylation of *h*20S

Commercially available *h*20S was dialyzed into PBS (Gibco Life Technologies) pH 7.5 and biotinylated with 20 eq of NHS-Biotin (Thermo Scientific, 20217) at 4 °C for 2 h. Excess reagent was removed with a PD-10 desalting column (Amersham Biosciences), separated into aliquots, snap-frozen, and stored at -80 °C for later use.

### Binding kinetics analysis

Biolayer interferometry (BLI) data of PA26 activators were measured using an Octet RED384 (ForteBio). The reactions were carried out in black 384-well plates (Greiner Bio-One, 781209) at 25 °C with a volume of 85 μl per well in BLI buffer (assay buffer containing 0.2% (w/v) BSA (Sigma)). Biotinylated *h*20S (10 nM) were immobilized on streptavidin (SA) biosensor. Serial dilutions of PA26 in BLI buffer were used as analyte. Association was observed by immersing loaded biosensors into solutions of analyte for 450 s. No binding was detected of the analyte to unloaded sensors. Dissociation was observed by transferring the sensor to a well containing binding buffer and no analyte for 1200 s. Affinity (*K*_D_) and kinetic parameters (*k*_on_ and *k*_off_) were calculated from a global fit (2:1 heterogenous ligand model) of the data using the Octet data analysis software.

### Cryo–electron microscopy grid preparation

*h*20S proteasome and PA26^E102A-Opt5^ activators were mixed by dilution into a buffer system (20 mM Tris pH 7.5, 20 mM NaCl, 10 mM KCl, 1 mM DTT, and 0.025% Nonidet P40) to a final concentration of 2 μM and 4 μM, respectively. The mixture was centrifuged at 25 °C for 30 min at 12,000 x g in a tabletop centrifuge and placed on ice.

Three microliters of the *h*20S-PA26^E102A-Opt5^ solution were applied to R2/2 400-mesh grids (Quantifoil) that had been plasma treated for 25 s using a glow discharger (Electron Microscopy Sciences) operated under atmospheric gases doped with amylamine. The grids were manually blotted to near dryness with Whatman no. 1 filter paper inside a cold room (4 °C, 95% humidity). The sample application and blotting process was repeated twice more to increase on-grid protein concentration. After the third blot, the grid was gravity plunged into liquid ethane using a home-made system and stored under liquid nitrogen.

### Cryo-electron microscopy data collection and image processing

Cryo-EM data were acquired using the Leginon software for automated data acquisition^54^ using a Titan Krios (Thermo Fisher) equipped with a K2 Summit (Gatan) direct electron detector in counting mode (**Supplementary Table 2**). Movies were collected by navigating to the center of a hole and image shifting a beam of ∼600 nm diameter to 10 targets situated at the periphery of the 2-μm hole. A total of 13,329 movies were recorded at a nominal magnification of 29,000 × (1.03 Å magnified pixel size) and composed of 29 frames (250 ms per frame, ∼50 e^−^/Å^2^ per movie). Movie collection was guided by real-time assessment of image quality using the Appion image processing environment^55^. Frame alignment and dose weighting were performed in real-time using UCSF Motioncor2^56^. CTF estimation on aligned, unweighted, micrographs was performed with Gctf^57^. All data post-processing steps were conducted in RELION 2.1^52,53^.

Single particle analysis was performed in RELION 2.1. Particle picking was conducted using the AutoPick function and resulted in 2,650,143 particle picks that were extracted using a box size of 420 × 420 pixels, down-sampled to 70 × 70 pixels, for reference-free 2D classification. 497,630 particles belonging to the 2D classes demonstrating features characteristic of secondary structural elements were subjected to 3D refinement and subsequent 3D classification (k = 8). A 3D template of the PA26-bound archaeal proteasome (PDB ID: 1YAU) was lowpass-filtered to 60 Å and used to guide the initial 3D refinement and 3D classification. 288,515 particles corresponding to 3D classes without artefactual features were chosen for further data processing. To minimize the detrimental effects of pseudo-symmetry (C2) on resolution, the raw particles were C2 symmetry expanded, 3D refined, and a python script was used to determine the x- and y-shifts required to reposition the proteasome gate at the center of the particle box. The particles at the new center were re-extracted to serve as individual asymmetric units without down-sampling in a box of 256 × 256 pixels, resulting in a total of 577,030 particles. A round of reference-free 2D classification enabled us to remove the ends of *h*20S particles that lacked an activator molecule. This combined expansion and classification approach yielded 521,860 particles. We subjected these particles to 3D classification (k = 8) and selected 384,939 particles whose parent 3D classes resembled a fully assembled *h*20S-PA26^E102A-Opt5^ complex. Following removal of particle duplicates with a python script, 326,676 particles were CTF and beam tilt refined. Further 3D classification without alignment (k = 10) revealed two classes with an unresolved alpha subunit helix. We attributed this finding to rotational misalignment around the C7 pseudo-symmetry, longitudinal axis of the complex and discarded these particles. The remaining 247,362 particles were then sorted and pruned by z-score, yielding 234,960 particles for final 3D refinement (2.9 Å) (**Supplementary Figs. 6 and 7**).

### Atomic model building

The atomic model was built by using the *h*20S and PA26 (PDB IDs: 5A0Q and 1YAU, respectively) as templates and rigid body fitting each subunit with Chimera into the electron density map. Subunits for which there was no density as a result of the symmetry expansion and re-extraction processing approach, were removed from the template. The modified templates were subject to one cycle of morphing and simulated annealing in PHENIX, followed by a total of 5 real-space refinement macrocycles with atomic displacement parameters, secondary structure restraints, local grid searches, and global minimization. After PHENIX refinement, manual real-space refinement was performed in Coot. Multiple rounds of real-space refinement in PHENIX (five macro cycles, Ramachandran and rotamer restraints, no morphing, no simulated annealing) and Coot were performed to address geometric and steric discrepancies identified by the RCSB PDB validation server. To ensure atomic models were not overfit as a result of real-space refinement, map-to-model FSCs were calculated with PHENIX^58^ (**Supplementary Fig. 7b**). All images were generated using UCSF Chimera and ChimeraX.

## Data availability

Structural data are available in the Electron Microscopy Databank and the RCSB Protein Databank (EMDB ID: 22259 and PDB ID: 6XMJ). Additional data supporting the findings of this manuscript are available from the corresponding author upon reasonable request.

## Acknowledgements

This work is dedicated to L. L. Kiessling on the occasion of her 60^th^ birthday. This work was supported by grants from the Tau Consortium and NIH AG053619 (to J.E.G.), NIH AG061697 (to G.C.L.), and NIH/NIGMS R01GM083960 (to A.S.). Additional support included an NSF GRFP fellowship and an HHMI Gilliam fellowship (to K.A.O.-N.) and American Cancer Society fellowship (A.H.P.). All cryo-EM data were collected at the Scripps Research electron microscopy facility and we thank B. Anderson for his microscope support. We thank J.C. Ducom at the Scripps Research High Performance Computing for computational support. Computational analyses of EM data were performed using shared instrumentation funded by NIH S10OD021634 to G.C.L. The authors thank the laboratory of C.S. Craik (University of California San Francisco) for assistance with peptide synthesis, the laboratory of P. Coffino (The Rockefeller University) for the PA26 construct, as well as A.K. Mapp, M.A. Ravalin and L. Pope for helpful suggestions and comments on the manuscript.

## Author contributions

K.A.O.-N. and J.E.G. designed the studies and wrote the manuscript. All authors edited the manuscript. K.A.O.-N., S.K.M., and N.C. conducted biochemistry experiments, performed data analysis, and generated necessary reagents. A.H.P. performed cryo-EM sample preparation, data collection, and data processing. J.E.G., K.A.O.-N., A.S. and G.C.L provided funding.

