## Supplemental Figures for "The YΦ Motif Defines the Structure-Activity Relationships of Human 20S Proteasome Activators"

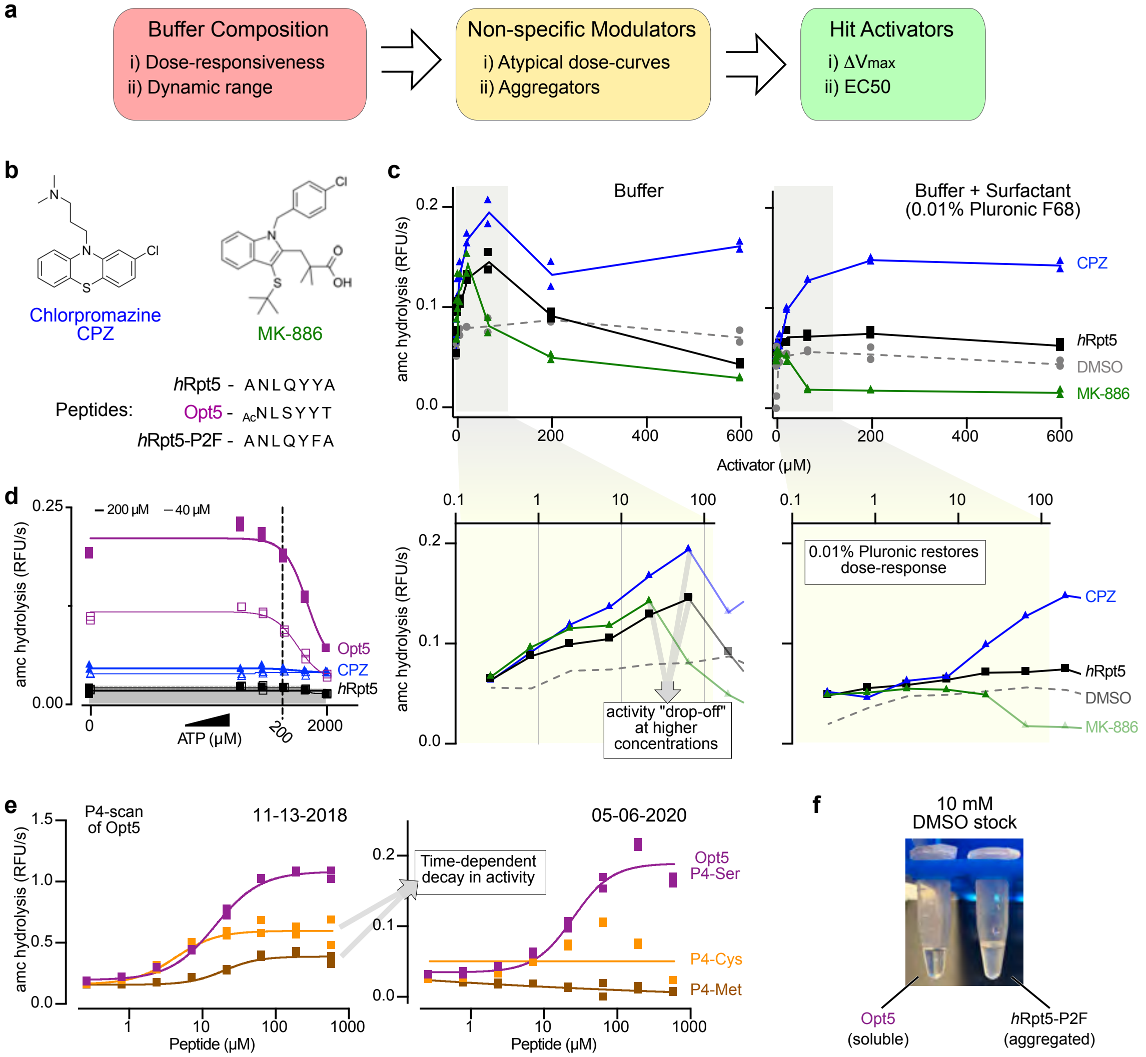

Supplementary Fig. 1. a, Pipeline for assay optimization and hit validation of peptide activators of the h20S. See text for details. Briefly, we started with a buffer previously reported to have optimized dynamic range, and then screened detergents to identify those that would not promote activity, but still minimize artifacts from aggregators. Finally, we measured both the effects on turnover rate (ΔVmax) and the EC50 (potency). b, Structures of 20S activators and peptide sequences used as benchmarks in this assay optimization study. c, Addition of surfactant (0.01% Pluronic F-68 ®) restores dose-dependence of h20S activation. Representative dose-response curves for small molecule (CPZ, blue; MK-886, green) and hRpt5 peptide (black) activators. Below each graph, a "zoom-in" is shown to highlight the non-specific inhibition at higher concentrations (above ~100 μM). We contribute this loss of activity to insolubility. Addition of detergent restores normal activity, except for the small molecule MK-886, which loses its activity (green). d, ATP inhibits activation by hRpt5-based peptides at high concentrations. Representative dose-response curve of activation with ATP titration. Based on these results, we chose an assay buffer that contains 200 μM of ATP. e, Time-dependent loss of activity in Cys- and Met-containing peptides. Two separate dose-response curves are shown on assays performed on different days (see Supplementary Fig. 4b). c-e, Technical replicates are plotted individually (n = 2). f, Typical, soluble peptide appears clear as 10 mM stock in DMSO (left, Opt5), while aggregated peptide appears white and cloudy (right, hRpt5-P2F).

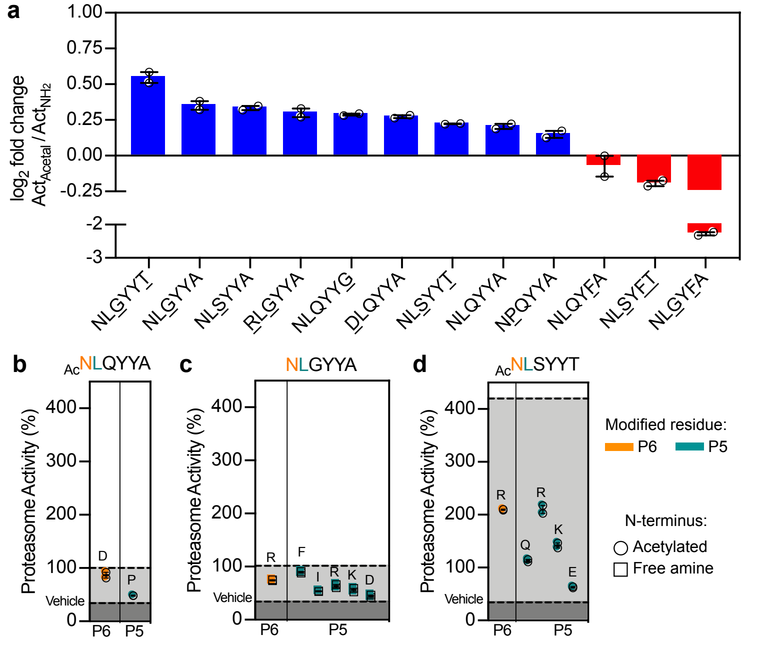

**Supplementary Fig. 2.** Effects of N-terminal modifications on stimulation of *h*20S by peptides. **a**, N-terminal acetylation of *h*Rpt5-derived peptides tends to modestly enhance their stimulatory activity. The effect of N-terminal acetylation is plotted as log2 fold increase (blue) or decrease (red) in proteasomal stimulation. Additional modifications from the wild type *h*Rpt5 sequence are underlined. Reported data is a mean calculated from two independent experiments with error reported as s.e.m. **b-d**, Scatterplots of the relative activities of peptides (250 μM) sampling residues at the P6 (orange) and P5 (teal) positions. Activities are benchmarked by vehicle and either (**b**) Ac-NLQYYA, (**c**) NLGYYA, or (**d**) Ac-NLSYYT (dotted lines). Data are normalized to Ac-NLQYYA and plotted individually (n = 2) along with the mean ± s.e.m.

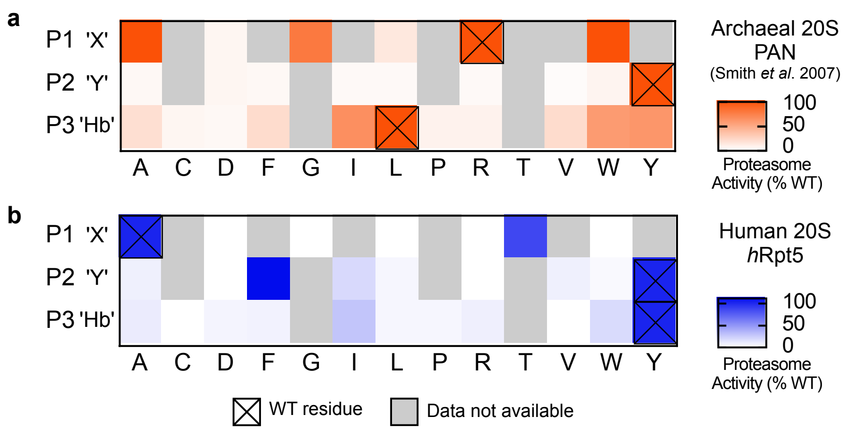

**Supplementary Fig. 3.** The structure-activity profile for *h*Rpt5-activated *h*20S differs from the HbYX model **a**, Summary of the previously described SAR for PAN-stimulated activity of the Thermoplasma 20S proteasome (9) (orange). **b**, The SAR for *h*Rpt5-stimulated activity of the human 20S (blue). The corresponding mutations from Smith et al. 2007 were made to Ac-NLQYYA and the data aligned to the archaeal heatmaps. Only a subset of the h20S results are included to allow direct comparisons to exact sequences. Data are normalized to WT and plotted as mean (n = 2). **a**,**b**, Residues of wildtype sequences are denoted with an ‘x’. Activity measurements were not determined for residues in gray.

**a**

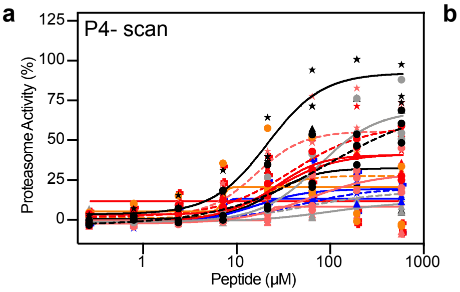

**b**

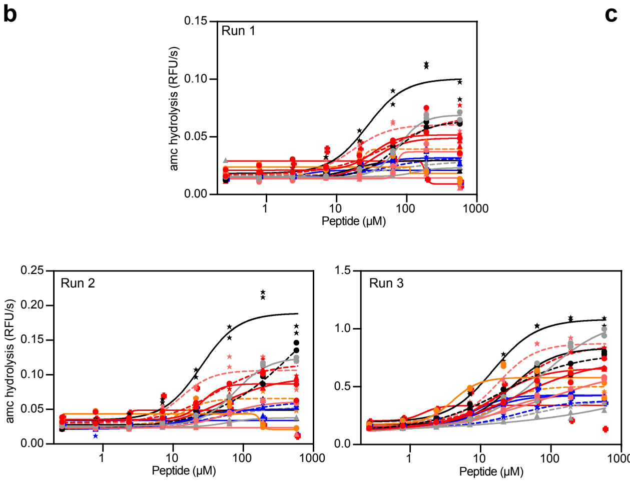

**c**

| P4 Scan | Run 1 | Run 2 | Run 3 | Mean (Raw) | Mean (Normalized) |
|---|---|---|---|---|---|
| | EC50 ± s.e.m. (μM) | | | | |
| $_{Ac}$NLAYYT | 33.2 ± 1.6 | 26.1 ± 1.3 | 43.0 ± 2.1 | 34.1 ± 4.9 | 27.9 ± 1.3 |
| $_{Ac}$NLDYYT | 77.1 ± 1.1 | 76.3 ± 1.1 | 76.9 ± 1.5 | 76.8 ± 0.2 | 74.9 ± 1.4 |
| $_{Ac}$NLEYYT | N.D. | 53.5 ± 1.2 | 83.6 ± 2.2 | 68.5 ± 15.0 | 53.1 ± 3.2 |
| $_{Ac}$NLFYYT$^{\Delta}$ | > 600 | > 600 | > 600 | N.D. | > 600 |
| $_{Ac}$NLGYYT | 68.7 ± 1.1 | > 600 | 20.5 ± 1.3 | 44.6 ± 24.1 | 44.6 ± 2.2 |
| $_{Ac}$NLHYYT | 33.9 ± 1.2 | 48.1 ± 1.1 | 15.9 ± 1.1 | 32.6 ± 9.3 | 25.8 ± 1.5 |
| $_{Ac}$NLIYYT $^{\Delta}$ | > 600 | > 600 | > 600 | N.D. | > 600 |
| $_{Ac}$NLKYYT | > 600 | > 600 | > 600 | N.D. | > 600 |
| $_{Ac}$NLLYYT | > 600 | > 600 | > 600 | N.D. | > 600 |
| $_{Ac}$NLNYYT | 17.1 ± 1.7 | 21.7 ± 1.4 | 27.9 ± 1.1 | 22.2 ± 3.1 | 23.8 ± 2.2 |
| $_{Ac}$NLPYYT | 62.3 ± 1.6 | 39.4 ± 1.3 | 29.8 ± 1.0 | 43.8 ± 9.6 | 39.2 ± 1.5 |
| $_{Ac}$NLQYYT | 53.6 ± 1.4 | 63.9 ± 2.0 | 32.9 ± 1.1 | 50.1 ± 9.1 | 46.9 ± 1.5 |
| $_{Ac}$NLRYYT | > 600 | > 600 | > 600 | N.D. | > 600 |
| $_{Ac}$NLSYYT | 27.5 ± 1.2 | 25.0 ± 1.2 | 15.0 ± 1.1 | 22.5 ± 3.8 | 21.4 ± 1.2 |
| $_{Ac}$NLTYYT | 17.0 ± 1.4 | 13.1 ± 1.4 | 18.0 ± 1.2 | 16.0 ± 1.5 | 16.2 ± 1.4 |
| $_{Ac}$NLVYYT | N.D. | 14.7 ± 1.5 | 18.0 ± 1.2 | 16.3 ± 1.6 | 16.7 ± 1.6 |
| $_{Ac}$NLWYYT$^{\Delta}$ | > 600 | > 600 | > 600 | N.D. | > 600 |
| $_{Ac}$NLYYYT $^{\Delta}$ | > 600 | > 600 | > 600 | N.D. | > 600 |

Δ = Positive for colloidal aggregates

**d**

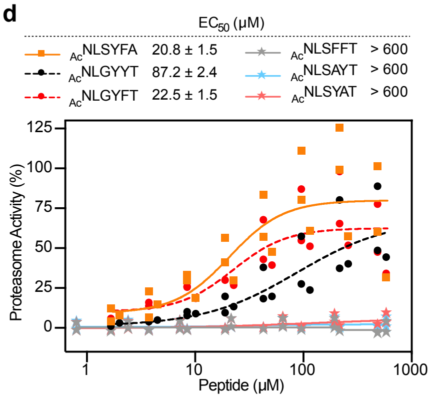

**e**

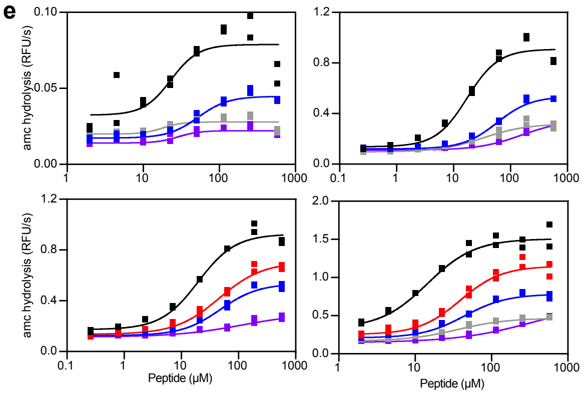

**f**

| | Run 1 | Run 2 | Run 3 | Run 4 | Mean (Raw) | Mean (Normalized) |
|---|---|---|---|---|---|---|
| | EC50 ± s.e.m. (μM) | | | | | |
| NLSYFT | 23.2 ± 1.5 | 17.3 ± 1.2 | 19.7 ± 1.1 | 14.5 ± 1.2 | 18.7 ± 1.8 | 15.9 ± 1.2 |
| NLSY2T | - | - | 46.1 ± 1.2 | 39.0 ± 1.1 | 42.5 ± 3.5 | 40.9 ± 1.1 |
| NLSY3T | 52.9 ± 1.1 | 58.3 ± 1.1 | 48.0 ± 1.2 | 44.2 ± 1.1 | 50.8 ± 3.1 | 51.0 ± 1.1 |
| NLSY4T | > 600 | > 600 | - | > 600 | N.D. | > 600 |
| NLSY5T | > 600 | > 600 | > 600 | > 600 | N.D. | > 600 |

**g**

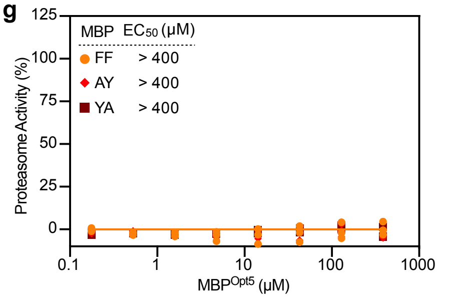

**h**

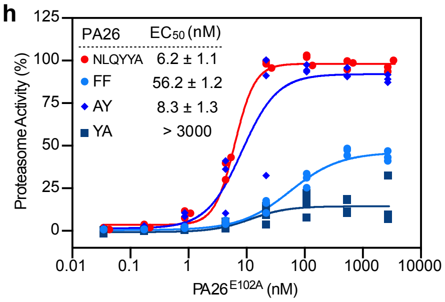

**Supplementary Fig. 4.** Dose-dependent stimulation induced by different PAs. **a-c**, Stimulation of *h*20S by P4-substituted hexapeptides. (**a**) Normalized dose-response curves. Data are normalized to Opt5 (Ac-NLSYYT) and plotted individually (n = 3). (**b**) Experimental replicates of non-normalized dose-response curves with technical replicate plotted individually (n=2). (**c**) EC50 values are reported for each independent experiment (n = 3). The mean of the "raw" EC50 values (yellow) along with the mean of the normalized EC50 values (blue) are reported with error as s.e.m. (n = 3). **d**. Dose-response curves for additional *h*Rpt5-derived hexapeptides. Data are normalized to Opt5 and plotted individually (n = 3) with calculated EC50 values with error reported as s.e.m. (n = 3). **e,f**, Activity of hexapeptides for pi-stacking studies. (**e**) Experimental replicates of raw dose-response curves with technical replicate plotted individually (n=2). (**f**) EC50 values are reported for each independent experiment (n = 4). The mean of the "raw" EC50 values (yellow) along with the mean of the normalized EC50 values (blue) are reported with error as s.e.m. (n = 2 to 4). **g,h.** Dose-response curves for additional (**g**) C-terminal MBP mutants and (**h**) C-terminal PA26E102A mutants. Data are normalized to Opt5 sequence and plotted individually (n = 3). Reported EC50 is a mean of EC50 values calculated from two to four independent experiments with error reported as s.e.m.

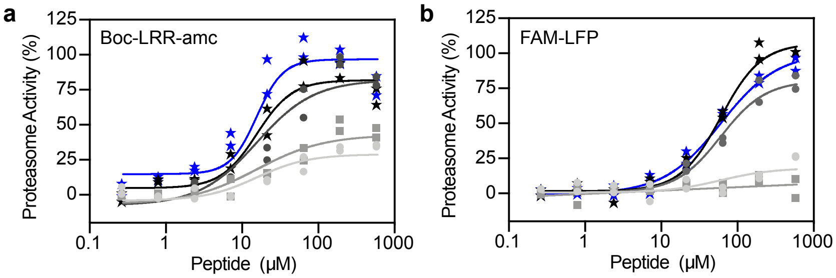

**c**

| $EC_{50}$ ± s.e.m. (μM) | Peptidic 20S Substrates LLVY | LRR | LFP |
|---|---|---|---|
| $_{Ac}$NLSYFT | 17.3 ± 1.1 | 9.3 ± 1.2 | 48.5 ± 1.1 |
| $_{Ac}$NLSYYT | 35.5 ± 1.1 | 15.3 ± 1.3 | 58.5 ± 1.1 |
| $_{Ac}$NLSYYA | 65.0 ± 1.1 | 35.1 ± 1.2 | 72.2 ± 1.2 |
| $_{Ac}$NLQYYT | 66.0 ± 1.4 | 42.2 ± 1.3 | > 600 |
| $_{Ac}$NLQYYA | > 600 | 47.3 ± 1.2 | > 600 |

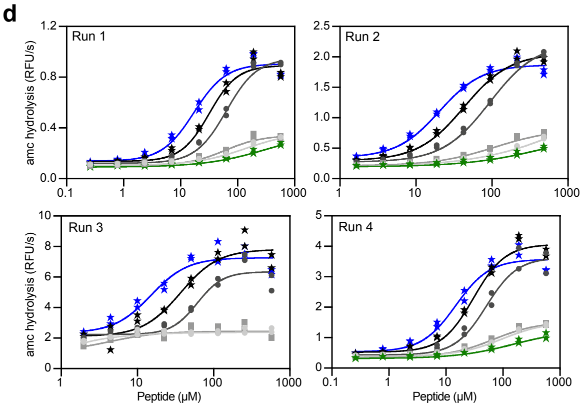

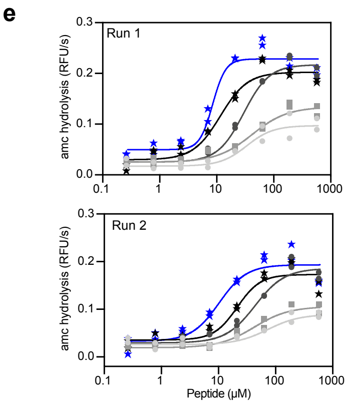

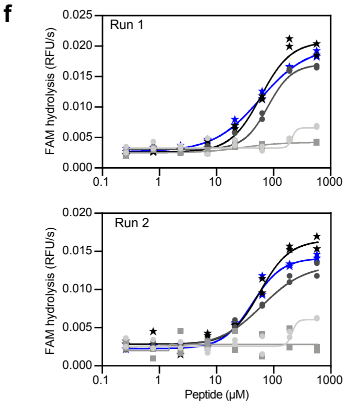

| LLVY | $EC_{50}$ ± s.e.m. (μM) Run 1 | Run 2 | Run 3 | Run 4 | Mean (Raw) | Mean (Normalized) |
|---|---|---|---|---|---|---|
| $_{Ac}$NLSYFT | 17.2 ± 1.2 | 20.4 ± 1.1 | 15.1 ± 1.3 | 14.9 ± 1.2 | 16.9 ± 1.3 | 17.3 ± 1.1 |
| $_{Ac}$NLSYYT | 31.9 ± 1.2 | 43.0 ± 1.1 | 36.8 ± 1.2 | 30.4 ± 1.1 | 35.5 ± 2.8 | 35.5 ± 1.1 |
| $_{Ac}$NLSYYA | 63.7 ± 1.1 | 93.8 ± 1.1 | 61.0 ± 1.2 | 51.1 ± 1.2 | 67.4 ± 9.2 | 65.0 ± 1.1 |
| $_{Ac}$NLQYYT | 60.5 ± 1.2 | 108 ± 1 | > 600 | 68.0 ± 1.2 | 78.8 ± 14.7 | 66.0 ± 1.4 |
| $_{Ac}$NLQYYA | > 600 | > 500 | > 600 | 96.1 ± 1.1 | N.D. | > 600 |
| $_{Ac}$NLSFYT | > 600 | > 500 | - | > 600 | N.D. | > 600 |

| LRR | $EC_{50}$ ± s.e.m. (μM) Run 1 | Run 2 | Mean (Raw) | Mean (Normalized) |
|---|---|---|---|---|
| $_{Ac}$NLSYFT | 10.3 ± 1.3 | 8.5 ± 1.2 | 9.4 ± 0.8 | 9.3 ± 1.2 |
| $_{Ac}$NLSYYT | 21.3 ± 1.2 | 11.6 ± 1.2 | 16.5 ± 4.8 | 15.3 ± 1.3 |
| $_{Ac}$NLSYYA | 43.8 ± 1.2 | 29.6 ± 1.1 | 36.7 ± 7.1 | 35.1 ± 1.2 |
| $_{Ac}$NLQYYT | 42.5 ± 1.3 | 41.8 ± 1.2 | 42.1 ± 0.4 | 42.2 ± 1.3 |
| $_{Ac}$NLQYYA | 73.2 ± 1.4 | 33.6 ± 1.4 | 53.4 ± 20 | 47.3 ± 1.2 |

| LFP | $EC_{50}$ ± s.e.m. (μM) Run 1 | Run 2 | Mean (Raw) | Mean (Normalized) |
|---|---|---|---|---|
| $_{Ac}$NLSYFT | 58.3 ± 1.1 | 42.5 ± 1.1 | 50.4 ± 7.9 | 48.5 ± 1.1 |
| $_{Ac}$NLSYYT | 59.5 ± 1.1 | 57.3 ± 1.1 | 58.4 ± 1.1 | 58.5 ± 1.1 |
| $_{Ac}$NLSYYA | 79.6 ± 1.1 | 62.4 ± 1.2 | 71.0 ± 8.6 | 72.2 ± 1.2 |
| $_{Ac}$NLQYYT | > 600 | > 600 | N.D. | > 600 |
| $_{Ac}$NLQYYA | > 600 | > 600 | N.D. | > 600 |

**Supplementary Fig. 5.** Distinct proteasome substrates report similar SAR for *h*Rpt5-based activators. **a**,**b**, Stimulation of the *h*20S (4 nM) by *h*Rpt5-based peptides was assessed with (**a**) the trypsin-targeted boc-LRR-amc (20 μM) and (**b**) the gate-sensitive nonapeptide FAM-LFP (100 nM) as alternate substrates to suc-LLVY-amc. Data are normalized to (**a**) Ac-NLSYFT or (**b**) Ac-NLSYYT and plotted individually (n = 2). **c**, **d** EC50 values determined with LRR and LFP substrates were calculated from two independent experiments and reported as the mean ± s.e.m. Corresponding values for LLVY were also included. **d**-**f**, Experimental replicates of non-normalized dose-response curves for optimization panel of *h*Rpt5 assessing hydrolysis of (**d**) LLVY, (**e**) LRR, and (**f**) LFP substrates. Technical replicates (n=2) are plotted individually for two to four independent experiments with the EC50 values ± s.e.m. reported for each replicate. The “raw” mean (yellow) EC50 was calculated and reported alongside the “normalized” mean (blue) with error reported as s.e.m. (n = 2 or 4).

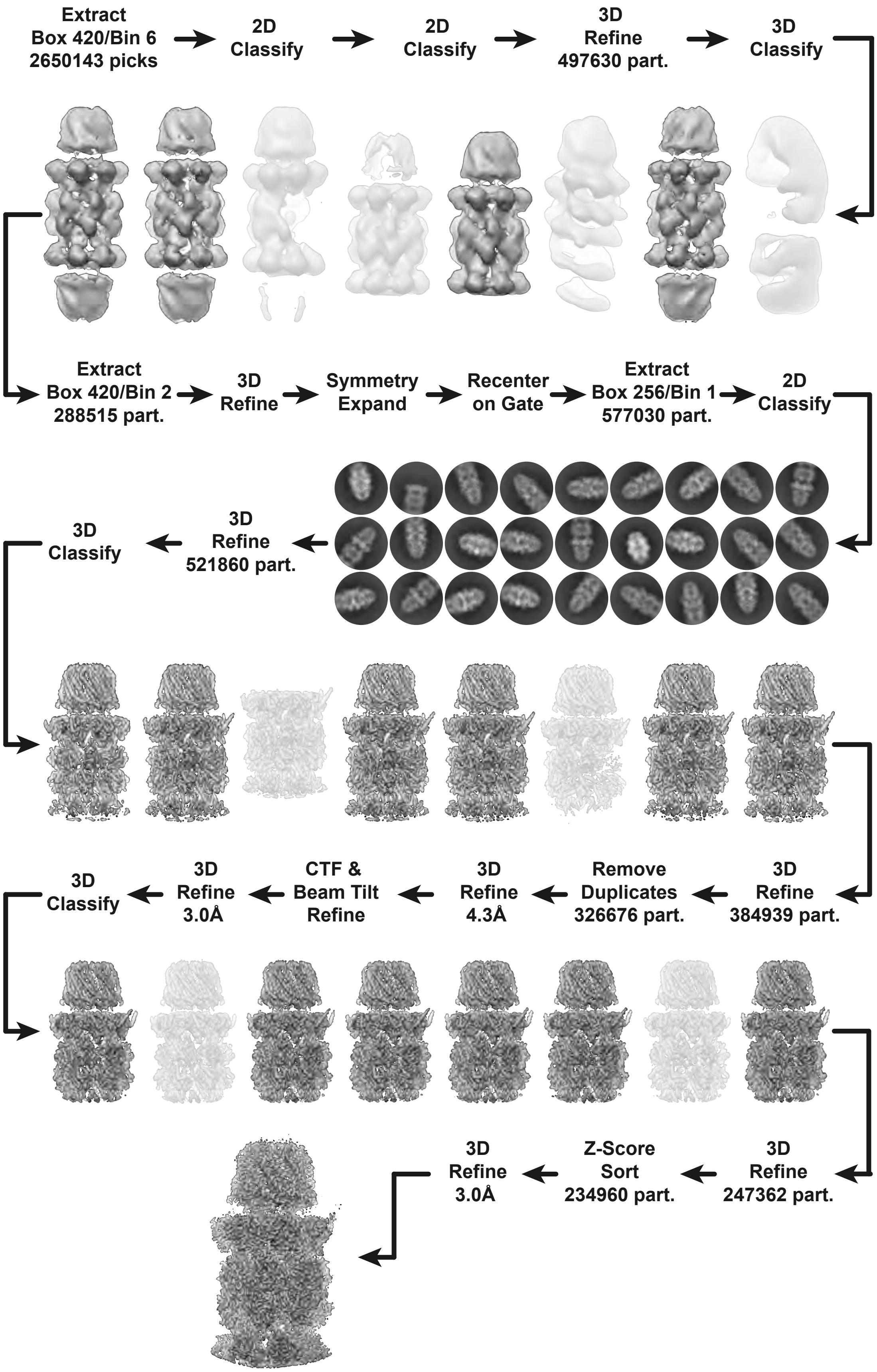

**Supplementary Fig. 6.** Schematic for cryo-EM single-particle data processing. Low resolution or artefactual reconstructions (white) and corresponding particles were excluded from subsequent processing steps, whereas all other reconstructions (gray) and corresponding particles were utilized. Representative 2D classes (dark circular background) are shown.

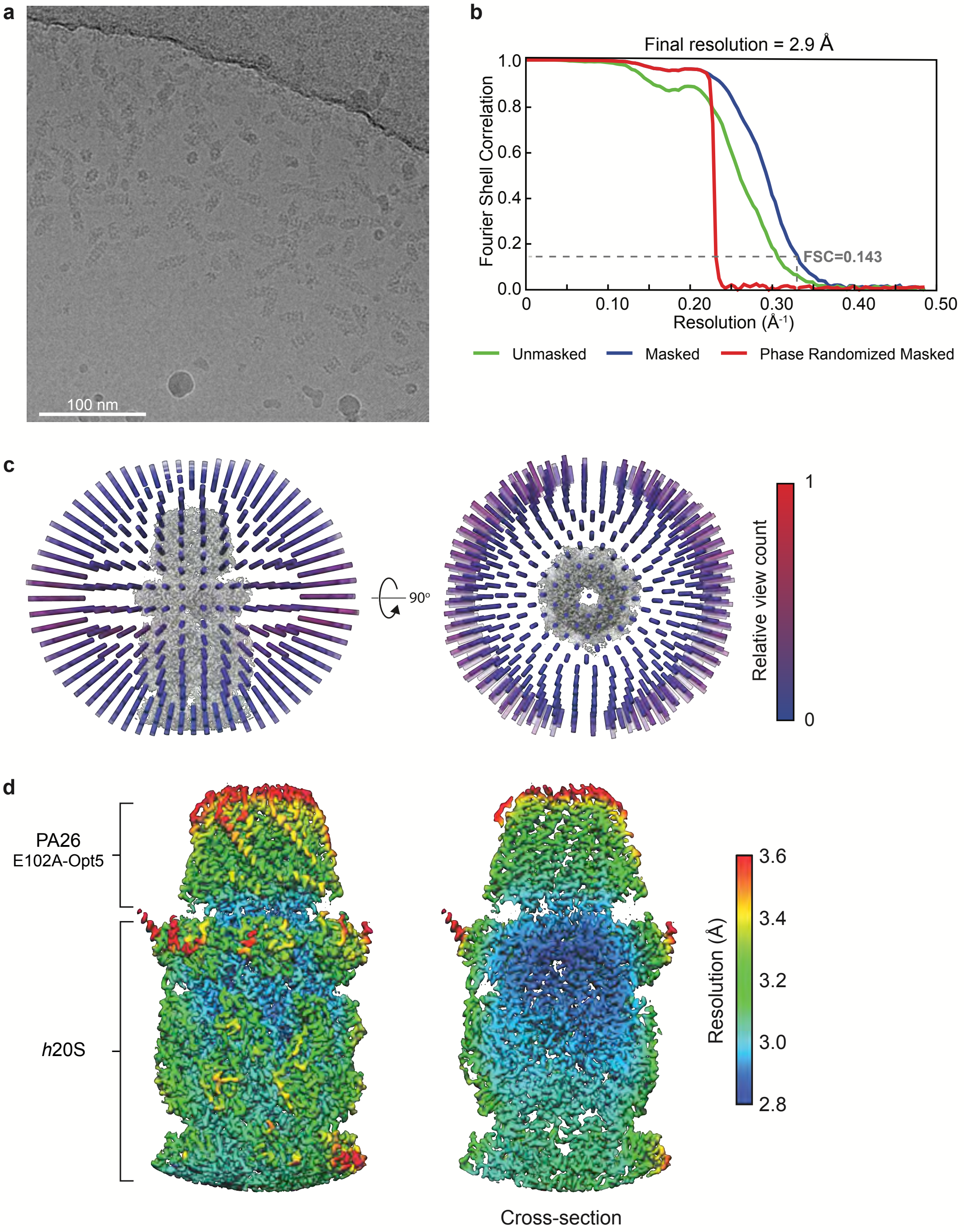

**Supplementary Fig. 7.** Cryo-EM metrics for PA26E102A-Opt5-*h*20S complex. **a**, Representative micrograph. **b**, The Fourier Shell Correlation (FSC) curves for the unmasked (green), masked (blue) and phase randomized (red) reconstructions. The resolution at 0.143 is indicated by a dashed line. **c**, Distribution of Euler angles displaying the orthogonal view of the reconstruction. The angular distribution is shown as a column whose longitudinal axis aligns with the normal of the corresponding back-projection **d**, Local resolution estimates of the global reconstruction for the complete volume (left) and a coronal cross-section (right). **c,d** Generated and calculated with RELION44,45.

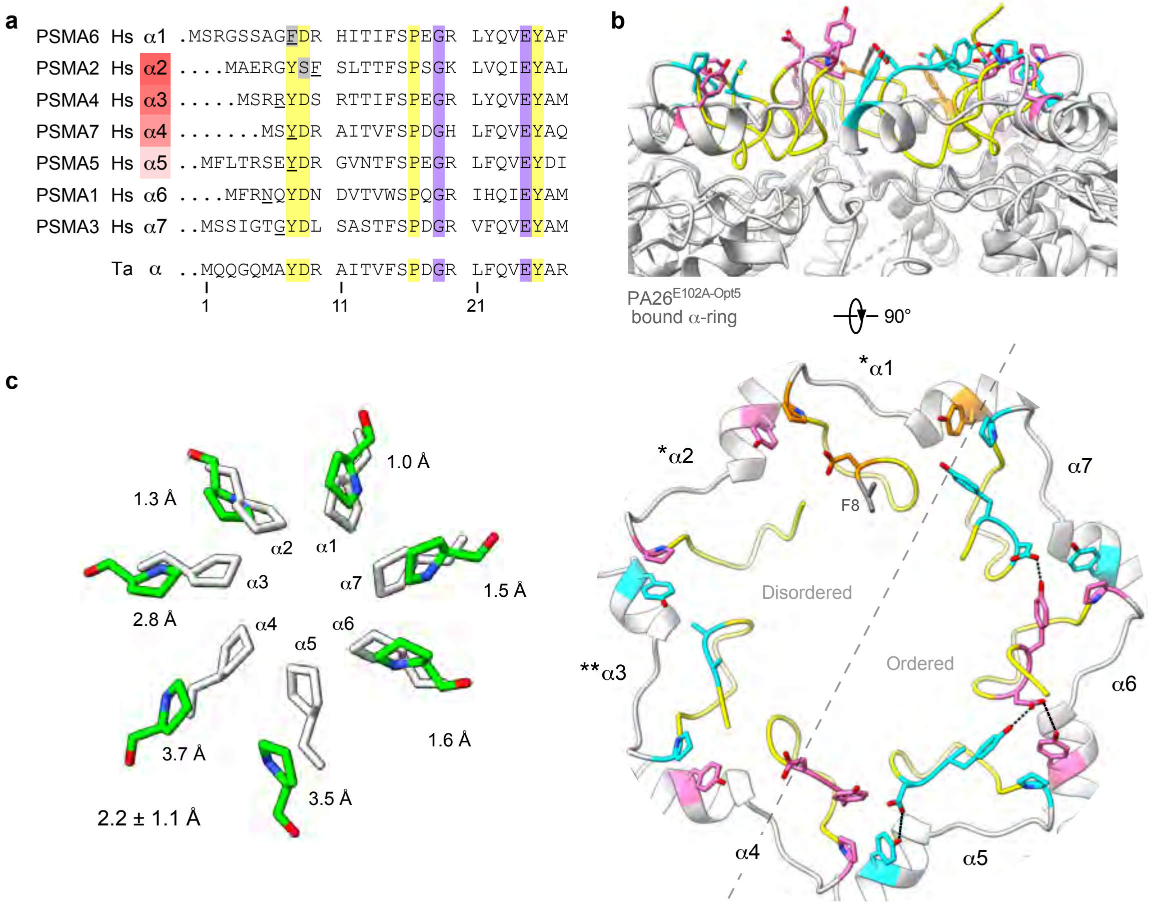

**Supplementary Fig. 8.** Asymmetric *h*20S open-gate conformation. **a**, N-terminal sequence alignment of 20S proteasome -subunits from *H. sapien* (Hs) and *T. acidophilum* (Ta). The α-subunits are labeled with human gene name but numbered according to yeast α-subunit/yeast gene. The α-subunits that directly gate the α-annulus (red) with their relative contributions to gating denoted with a gradient. The most N-terminal residue with resolvable backbone density in the cryo-EM reconstruction is underlined. The conserved cluster (yellow), non-canonical residues (gray), and conserved α-pocket residues (purple) are highlighted. **b**, Conformational state of N-terminal gate (yellow) and interactions of conserved cluster (α1, orange; α2, α4, α6, pink; α3, α5, α7, cyan) that stabilize asymmetric open-gate *h*20S. Non-canonical residues (gray) are denoted for resolved residues. Single or double asterisks indicates α-subunits featuring unresolved extreme N-termini with non-canonical (*) or conserved (**) residues, respectively. **c**, Radial displacement of Pro17. Individual measurements are reported along with mean ± s.d.

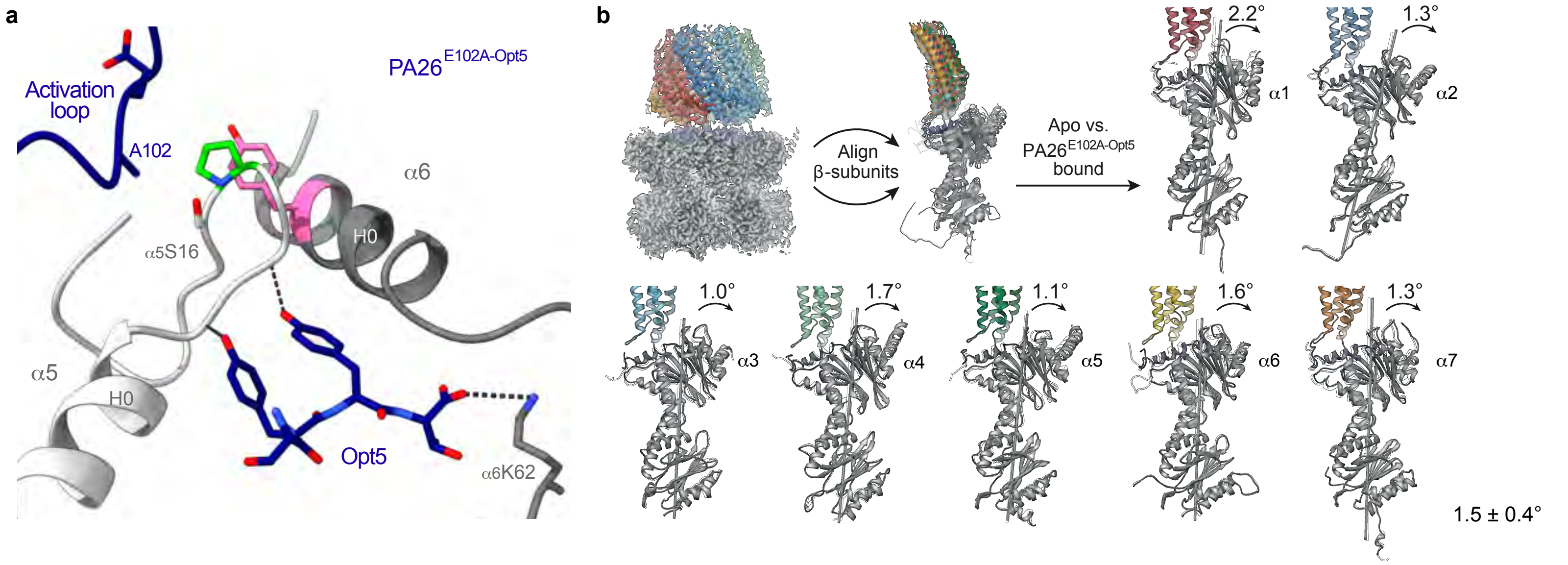

**Supplementary Fig. 9.** Structural evidence implicates termini-induced gate opening**. a**, Disabled activation loop of PA26E102A-Opt5 no longer interacts with αPro17 reverse turn in representative α-pocket (α5/α6-pocket). C-terminal Opt5 engages residues within reverse-turn loop from the α-pocket. **b**, PA26E102A-Opt5 induces rotation of the α-subunits about the axial channel. Individual measurements are reported along with mean ± s.d.

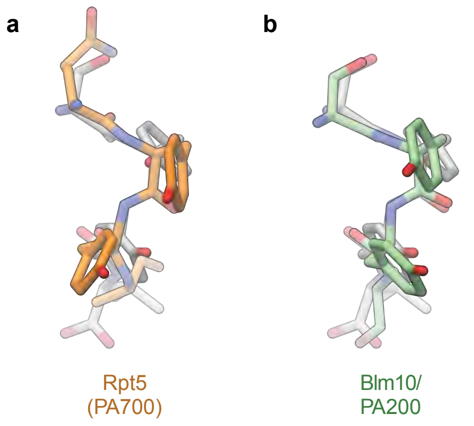

**Supplementary Fig. 10**. Binding orientation of reported YΦ motifs align with those observed for Opt5. **a,b,** Overlay highlights similar backbone conformation between bound Opt5 (SYYT, grey) and the C-terminus of (**a**) the human Rpt5 (QYYA, orange) (PDB ID: 5GJR) or (**b**) the yeast Blm10/PA200 (SYYA, green) (PDB ID: 4V70). Note that the aromatic P2 and P3 Tyr residues engage in off-set π-stacking.

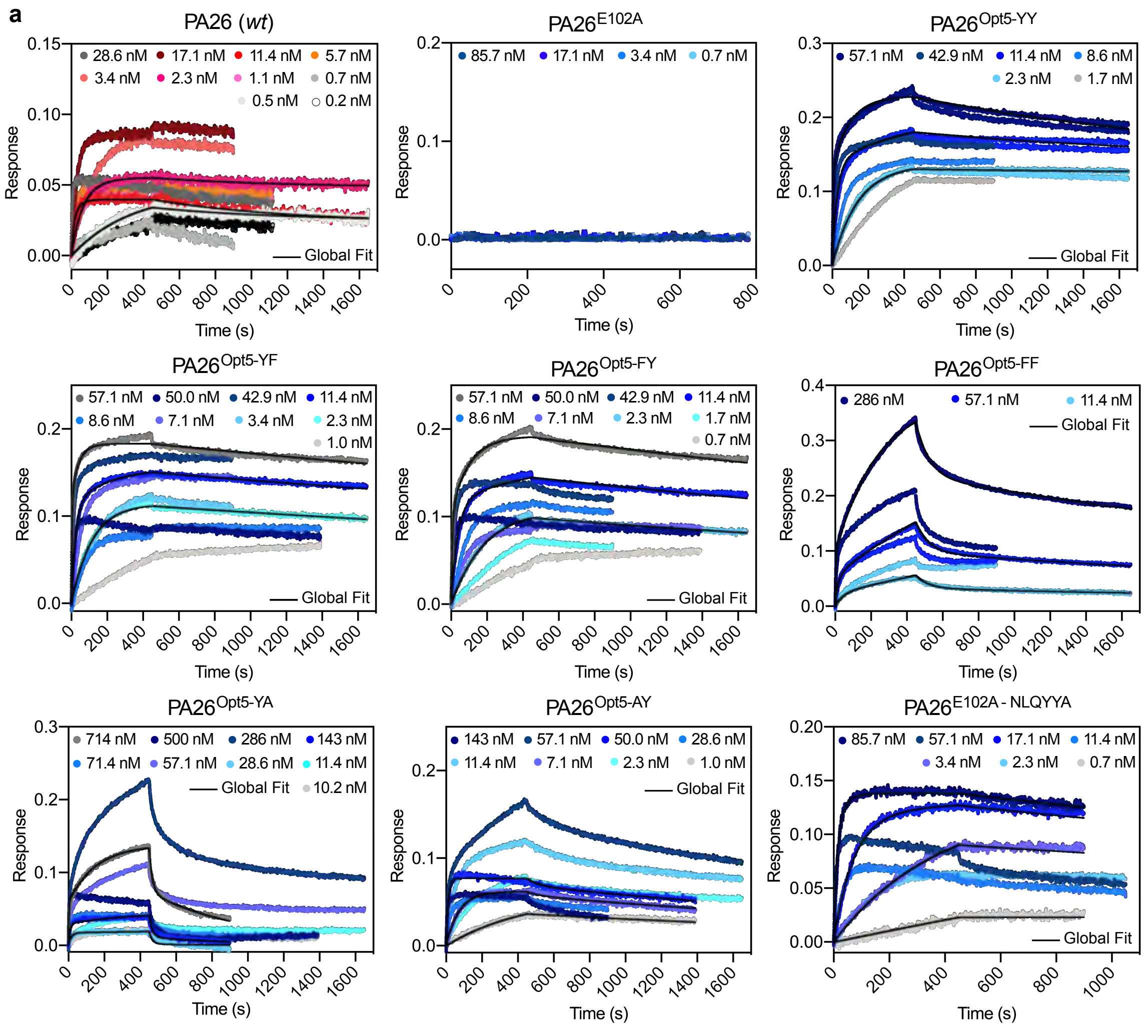

**b**

| PA26 | $K_D$ ± s.e.m. ($10^{-11}$ M) | $k_{on}$ ± s.e.m. ($10^4$ $M^{-1}s^{-1}$) | $k_{off}$ ± s.e.m. ($10^{-4}$ $s^{-1}$) |
|---|---|---|---|
| *wt* | 10 ± 5 | 710±310 | 2.9±0.8 |
| E102A | > 9000 | N.D. | N.D. |
| Opt5-YY NLSYYT | 36 ± 16 | 49±4 | 1.6±0.7 |
| Opt5-YF NLSYFT | 22 ± 10 | 150±30 | 2.6±0.5 |
| Opt5-FY NLSFYT | 38 ± 16 | 120±60 | 3.5±1.6 |
| Opt5-FF NLSFFT | 3500±1800 | 1.2±0.3 | 79±75 |
| Opt5-YA NLSYAT | 3600±900 | 27±26 | 110±110 |
| Opt5-AY NLSAYT | 28±13 | 150±20 | 4.0±1.6 |
| E102A NLQYYA | 9.7±2.9 | 190±110 | 1.5±0.5 |

**Supplementary Fig. 11**. Biolayer interferometry (BLI) of PA26 activators. **a**, Sensorgram traces of *h*20S binding to either wildtype (*wt*, red) or E102A (blue) variants of PA26 at different concentrations. Colored curves represent measurements for individual experiments (n = 2 or 3) and solid black lines correspond to global-fit for a representative experiment. **b,** Binding and kinetic constants measured by BLI for binding of PA26 to *h*20S. N.D. denotes values that could not be determined due to undetectable binding.

**Supplementary Table 1.** *h*Rpt5-derived peptides

| ID | Peptide length | N-term Modification | Sequence | Modification |
|---|---|---|---|---|
| 1 | 7 | - | ANLQYYA | - |
| 2 | 7 | - | AALQYYA | NP6A |
| 3 | 7 | - | ANAQYYA | LP5A |
| 4 | 7 | - | ANLAYYA | QP4A |
| 5 | 7 | - | ANLQAYA | YP3A |
| 6 | 7 | - | ANLQYAA | YP2A |
| 7 | 6 | - | NLQYYA | - |
| 8 | 5 | - | LQYYA | - |
| 9 | 4 | - | QYYA | - |
| 10 | 6 | Ac | NLQYYA | - |
| 11 | 6 | Ac | NLQYYA-CONH$_2$ | C-term amide |
| 12 | 6 | - | NLQYYD | AP1D |
| 13 | 6 | Ac | NLQYYD | AP1D |
| 14 | 6 | - | NLQYYG | AP1G |
| 15 | 6 | Ac | NLQYYG | AP1G |
| 16 | 6 | Ac | NLQYYL | AP1L |
| 17 | 6 | Ac | NLQYYR | AP1R |
| 18 | 6 | Ac | NLQYYT | AP1T |
| 19 | 6 | Ac | NLQYYW | AP1W |
| 20 | 6 | Ac | NLQYAA | YP2A |
| 21 | 6 | Ac | NLQYDA | YP2D |
| 22 | 6 | - | NLQYFA | YP2F |
| 23 | 6 | Ac | NLQYFA | YP2F |
| 24 | 6 | Ac | NLQYIA | YP2I |
| 25 | 6 | Ac | NLQYLA | YP2L |
| 26 | 6 | Ac | NLQYRA | YP2R |
| 27 | 6 | Ac | NLQYVA | YP2V |
| 28 | 6 | Ac | NLQYWA | YP2W |
| 29 | 6 | Ac | NLQAYA | YP3A |
| 30 | 6 | Ac | NLQCYA | YP3C |
| 31 | 6 | Ac | NLQDYA | YP3D |
| 32 | 6 | - | NLQFYA | YP3F |
| 33 | 6 | Ac | NLQFYA | YP3F |
| 34 | 6 | Ac | NLQIYA | YP3I |
| 35 | 6 | Ac | NLQLYA | YP3L |
| 36 | 6 | Ac | NLQPYA | YP3P |
| 37 | 6 | Ac | NLQRYA | YP3R |
| 38 | 6 | Ac | NLQSYA | YP3S |
| 39 | 6 | Ac | NLQVYA | YP3V |
| 40 | 6 | - | NLQWYA | YP3W |
| 41 | 6 | Ac | NLQWYA | YP3W |
| 42 | 6 | - | NLGYYA | QP4G |
| 43 | 6 | Ac | NLGYYA | QP4G |
| 44 | 6 | - | NLSYYA | QP4S |
| 45 | 6 | Ac | NLSYYA | QP4S |
| 46 | 6 | Ac | NLAYYT | AP1T, QP4A |
| 47 | 6 | Ac | NLCYYT | AP1T, QP4C |
| 48 | 6 | Ac | NLDYYT | AP1T, QP4D |
| 49 | 6 | Ac | NLEYYT | AP1T, QP4E |
| 50 | 6 | Ac | NLFYYT | AP1T, QP4F |
| 51 | 6 | - | NLGYYT | AP1T, QP4G |
| 52 | 6 | Ac | NLGYYT | AP1T, QP4G |
| 53 | 6 | Ac | NLHYYT | AP1T, QP4H |
| 54 | 6 | Ac | NLIYYT | AP1T, QP4I |
| 55 | 6 | Ac | NLKYYT | AP1T, QP4K |
| 56 | 6 | Ac | NLLYYT | AP1T, QP4L |
| 57 | 6 | Ac | NLMYYT | AP1T, QP4M |
| 58 | 6 | Ac | NLNYYT | AP1T, QP4N |
| 59 | 6 | Ac | NLPYYT | AP1T, QP4P |
| 60 | 6 | Ac | NLRYYT | AP1T, QP4R |
| 61 | 6 | - | NLSYYT | AP1T, QP4S |
| 62 | 6 | Ac | NLSYYT | AP1T, QP4S |
| 63 | 6 | Ac | NLTYYT | AP1T, QP4T |
| 64 | 6 | Ac | NLVYYT | AP1T, QP4V |

| | | | | |
|---|---|---|---|---|
| 65 | 6 | Ac | NLWYYT | AP1T, QP4W |
| 66 | 6 | Ac | NLYYYT | AP1T, QP4Y |
| 67 | 6 | - | RLGYYA | QP4G, NP6R |
| 68 | 6 | Ac | RLGYYA | QP4G, NP6R |
| 69 | 6 | - | DLQYYA | QP6D |
| 70 | 6 | Ac | DLQYYA | QP6D |
| 71 | 6 | - | NPQYYA | LP5P |
| 72 | 6 | Ac | NPQYYA | LP5P |
| 73 | 6 | - | NFGYYA | QP4G, LP5F |
| 74 | 6 | - | NIGYYA | QP4G, LP5I |
| 75 | 6 | - | NRGYYA | QP4G, LP5R |
| 76 | 6 | - | NKGYYA | QP4G, LP5K |
| 77 | 6 | - | NDGYYA | QP4G, LP5D |
| 78 | 6 | Ac | RLSYYT | AP1T, QP4S, NP6R |
| 79 | 6 | Ac | NQSYYT | AP1T, QP4S, LP5Q |
| 80 | 6 | Ac | NRSYYT | AP1T, QP4S, LP5R |
| 81 | 6 | Ac | NKSYYT | AP1T, QP4S, LP5K |
| 82 | 6 | Ac | NESYYT | AP1T, QP4S, LP5E |
| 83 | 6 | - | NLSFYA | YP3F, QP4S |
| 84 | 6 | Ac | NLSFYA | YP3F, QP4S |
| 85 | 6 | Ac | NLSYFA | YP2F, QP4S |
| 86 | 6 | - | NLGYFA | YP2F, QP4G |
| 87 | 6 | Ac | NLGYFA | YP2F, QP4G |
| 88 | 6 | - | NLGFYA | YP3F, QP4G |
| 89 | 6 | Ac | NLGFYA | YP3F, QP4G |
| 90 | 6 | - | NLGHYA | YP3H, QP4G |
| 91 | 6 | Ac | NLGHYA | YP3H, QP4G |
| 92 | 6 | - | NLSHYA | YP3H, QP4S |
| 93 | 6 | Ac | NLSHYA | YP3H, QP4S |
| 94 | 6 | Ac | NLGYFT | AP1T, YP2F, QP4G |
| 95 | 6 | - | NLSFYT | AP1T, YP3F, QP4S |
| 96 | 6 | Ac | NLSFYT | AP1T, YP3F, QP4S |
| 97 | 6 | - | NLSYFT | AP1T, YP2F, QP4S |
| 98 | 6 | Ac | NLSYFT | AP1T, YP2F, QP4S |
| 99 | 6 | - | NLSFFT | AP1T, YP2F, YP3F, QP4S |
| 100 | 6 | Ac | NLSFFT | AP1T, YP2F, YP3F, QP4S |
| 101 | 6 | - | NLSY4T | AP1T, YP2 4, QP4S |
| 102 | 6 | - | NLS4YT | AP1T, YP3 4, QP4S |
| 103 | 6 | - | NLSY2T | AP1T, YP2 2, QP4S |
| 104 | 6 | - | NLSY3T | AP1T, YP2 3, QP4S |
| 105 | 6 | - | NLSY5T | AP1T, YP2 5, QP4S |
| 106 | 6 | Ac | NLSYAT | AP1T, YP2A, QP4S |
| 107 | 6 | Ac | NLSAYT | AP1T, YP3A, QP4S |
| 108 | 6 | - | NLQYHA | YP2H |
| 109 | 6 | - | NLAYYA | QP4A |
| 110 | 6 | - | NLEYYA | QP4E |
| 111 | 6 | - | NLLYYA | QP4L |
| 112 | 6 | - | NLNYYA | QP4N |
| 113 | 6 | - | NLPYYA | QP4P |
| 114 | 6 | - | NLRYYA | QP4R |
| 115 | 6 | - | NLTYYA | QP4T |
| 116 | 6 | Ac | NLSYHT | AP1T, YP2H, QP4S |
| 117 | 6 | Ac | NLSYWT | AP1T, YP2W, QP4S |

**Supplementary Table 2:** Cryo-EM data collection, refinement, and validation statistics

| | |
|---|---|
| PDB ID | 6XMJ |
| EMDB ID | 22259 |
| **Data collection and Processing** | |
| Microscope | Titan Krios |
| Camera | K2 Summit |
| Magnification | 29,000 |
| Voltage (kV) | 300 keV |
| Total electron fluence ($e^{-}$/Å$^2$) | 50 |
| Electron flux ($e^{-}$/pixel/sec) | 8 |
| Defocus range (μm) | -1.5 to -3.0 |
| Pixel size (Å) | 1.03 |
| Micrographs collected (no.) | 13,392 |
| Total extracted particles (no.) | 2,650,143 |
| Refined particles (no.) | 497,630 |
| **Reconstruction** | |
| Final particles (no.) | 234,960 |
| Symmetry | C1 |
| Resolution (global, Å) | |
| FSC 0.5 | 3.1/3.0 |
| (unmasked/masked) | 3.0/2.9 |
| FSC 0.143 | 2.8-3.6 |
| (unmasked/masked) | |
| Resolution Range (local, Å) | |
| **Model composition** | |
| Nonhydrogen atoms | 64,832 |
| Protein residues | 4,555 |
| Ligands | 0 |
| Waters | 0 |
| **Refinement** | |
| MapCC (volume/masked) | 0.80/0.84 |
| Map sharpening *B* factor (Å$^2$) | -108 |
| R.m.s. deviations | |
| Bond lengths (Å) | 0.004 |
| Bond angles (°) | 0.746 |
| **Validation** | |
| EMRinger score | 3.97 |
| CaBLAM outliers (%) | 1.79 |
| MolProbity score | 1.13 |
| Clashscore | 1.77 |
| Rotamer outliers (%) | 0 |
| Ramachandran plot | |
| Outliers (%) | 0.02 |
| Allowed (%) | 3.19 |
| Favored (%) | 96.79 |

**Supplementary Table 3.** Measurements for *h*20S-PA26$^{\text{E102A-Opt5}}$ structure

α-subunit

| | α1 | α2 | α3 | α4 | α5 | α6 | α7 |
|---|---|---|---|---|---|---|---|
| Pro17 displacement (Å) | 1.0 | 1.3 | 2.8 | 3.7 | 3.5 | 1.6 | 1.5 |
| Rigid body rotation (°) | 2.2 | 1.3 | 1.0 | 1.7 | 1.1 | 1.6 | 1.3 |
| Unresolved gating residues (#) | 8 | 7 | 7 | 2 | 7 | 3 | 6 |

C-tail (α-pocket)

| | α7/α1 | α1/α2 | α2/α3 | α3/α4 | α4/α5 | α5/α6 | α6/α7 |
|---|---|---|---|---|---|---|---|
| C-tail electron density ($Å^3$) | 4.7 | 165.8 | 196.8 | 161.8 | 189.0 | 170.2 | 73.9 |
| Salt bridging distance (Å) | - | 5.1 | 3.4 | 3.2 | 2.9 | 3.6 | 4.3 |
| P2-P3 $\phi$ (°) | - | -61.5 | -127.3 | -121.1 | -137.8 | -131.8 | -129.9 |
| P2-P3 $\psi$ (°) | - | -39.8 | 159.2 | 167.2 | 149.2 | 136.3 | 150.9 |
| $\pi$-stacking R (Å) | - | 4.3 | 4.6 | 4.6 | 4.7 | 4.4 | 4.7 |
| $\pi$-stacking $\theta$ (°) | - | 25.7 | 26.0 | 26.3 | 23.7 | 33.3 | 25.6 |

α-ring pore

| | α1-α4 | α1-α5 | α2-α5 | α2-α6 | α3-α6 | α3-α7 | α4-α7 |
|---|---|---|---|---|---|---|---|
| Diameter (closed-gate) (Å)[a] | 37.5 | 39.2 | 37.7 | 36.9 | 36.4 | 36.8 | 36.9 |
| Diameter (open-gate) (Å) | 41.1 | 41.0 | 40.9 | 41.1 | 41.2 | 41.2 | 41.2 |

[a]Diameters are defined as the average distance between αPro17 Cα atoms across the α-ring denoted by the α-subunit pairs in the *h*20S (PDB ID: 4R3O)[32].

**Supplementary Table 4.** C-terminal sequences of PA200, PAN, Rpt5, Rpt3 and Rpt2 from different organisms identified in UniProtKB.

| Protein | Entry | Sequence | Entry name |
|---|---|---|---|
| PAN | Q58576 | LDVLYR | PAN_METJA |
| PAN | D4GUJ7 | VSRAFA | PAN1_HALVD |
| PAN | Q5UT56 | FTDYQY | PAN2_HALVD |
| PAN | Q9V2V6 | DEQTEE | PSA1_HALVD |
| PAN | D4GYZ1 | NFEGLE | PSB_HALVD |
| PAN | P9WQN5 | NLGQYL | ARC_MYCTU |
| PAN | Q9HRW6 | VSRTFA | PAN1_HALSA |
| PAN | Q9V287 | HEIIYG | PAN_PYRAB |
| PAN | O50202 | NTGQYL | ARC_RHOER |
| PAN | Q8TI88 | ETTMFV | PAN_METAC |
| PAN | Q8PY58 | PETMFV | PAN_METMA |
| PAN | Q9HNP9 | YPSYIQ | PAN2_HALSA |
| PAN | Q9V2V5 | TDEREE | PSA2_HALVD |
| PAN | O28303 | KGVMFV | PAN_ARCFU |
| PAN | Q8U4H3 | HEVIYG | PAN_PYRFU |
| PAN | O57940 | HEVIYG | PAN_PYRHO |
| PAN | Q5JHS5 | HEVMYG | PAN_THEKO |
| PAN | Q980M1 | RREKYS | PAN_SACS2 |
| PAN | C5A6P8 | HEVMYG | PAN_THEGJ |
| PAN | Q975U2 | RTEKYV | PAN_SULTO |
| PAN | Q6LWR0 | LTVMYG | PAN_METMP |
| PAN | A9A916 | LTVMYG | PAN_METM6 |
| PAN | Q2FQ56 | AGVMFA | PAN_METHJ |
| PAN | A6UQT3 | LTTMYG | PAN_METVS |
| PAN | O26824 | TGVMFG | PAN_METTH |
| PAN | A6VHR1 | LTVMYG | PAN_METM7 |
| PAN | B8GGN4 | SGAMFA | PAN_METPE |
| PAN | C3MY47 | RREKYS | PAN_SULIM |
| PAN | Q0W257 | SGVMFA | PAN_METAR |
| PAN | A7I8B8 | EGRMFA | PAN_METB6 |
| PAN | Q8TX03 | FKRAYH | PAN_METKA |
| PAN | C3N7K8 | RREKYS | PAN_SULIY |
| PAN | Q9YAC7 | TIATVI | PAN_AERPE |
| PAN | A4G0S4 | LTVMYG | PAN_METM5 |
| PAN | A3CV35 | FGEMFA | PAN_METMJ |
| PAN | C3MRF1 | RREKYS | PAN_SULIL |
| PAN | C3NFW6 | RREKYS | PAN_SULIN |
| PAN | C4KIR6 | RREKYS | PAN_SULIK |
| PAN | B6YXR2 | HEVMYG | PAN_THEON |
| PAN | C3MZI6 | RREKYS | PAN_SULIA |
| PAN | P9WQN4 | NLGQYL | ARC_MYCTO |
| PA200 | Q14997 | SPCYYA | PSME4_HUMAN |
| PA200 | Q5SSW2 | SPCYYA | PSME4_MOUSE |
| PA200 | F1MKX4 | SPCYYA | PSME4_BOVIN |
| PA200 | Q6NRP2 | SPCYYA | PSME4_XENLA |
| PA200 | F1QFR9 | SPCYYA | PSM4A_DANRE |
| PA200 | P43583 | WRSYYA | BLM10_YEAST |
| PA200 | F4JC97 | SSSYFA | PSME4_ARATH |
| Rpt5 | Q9SEI2 | SLNYYA | PS6AA_ARATH |
| Rpt5 | O76371 | SLNYYA | PRS6A_CAEEL |
| Rpt5 | Q54PN7 | TLEYYA | PRS6A_DICDI |
| Rpt5 | Q8SR13 | KLLYFT | PRS6A_ENCCU |
| Rpt5 | P17980 | NLQYYA | PRS6A_HUMAN |
| Rpt5 | O88685 | NLQYYA | PRS6A_MOUSE |
| Rpt5 | Q63569 | NLQYYA | PRS6A_RAT |
| Rpt5 | P33297 | SVSFYA | PRS6A_YEAST |
| Rpt5 | O42587 | NLQYYA | PR6AA_XENLA |

| Protein | Entry | Sequence | Entry name |
|---|---|---|---|
| Rpt3 | P43686 | EHEFYK | PRS6B_HUMAN |
| Rpt3 | P54775 | EHEFYK | PRS6B_MOUSE |
| Rpt3 | Q63570 | EHEFYK | PRS6B_RAT |
| Rpt3 | P33298 | KFDFYK | PRS6B_YEAST |
| Rpt3 | Q9SEI4 | DFEFYK | PRS6B_ARATH |
| Rpt3 | Q3T030 | EHEFYK | PRS6B_BOVIN |
| Rpt3 | O74894 | QFAFYK | PRS6B_SCHPO |
| Rpt3 | Q4R7L3 | EHEFYK | PRS6B_MACFA |
| Rpt3 | P46502 | DFEFYK | PRS6B_CAEEL |
| Rpt2 | P40327 | LEGLYL | PRS4_YEAST |
| Rpt2 | Q9SZD4 | PEGLYM | PRS4A_ARATH |
| Rpt2 | P62191 | PEGLYL | PRS4_HUMAN |
| Rpt2 | P62192 | PEGLYL | PRS4_MOUSE |
| Rpt2 | P48601 | PEGLYL | PRS4_DROME |
| Rpt2 | P62193 | PEGLYL | PRS4_RAT |
| Rpt2 | O16368 | PEELYL | PRS4_CAEEL |
| Rpt2 | Q90732 | PEGLYL | PRS4_CHICK |
| Rpt2 | Q8SRH0 | SAGLYS | PRS4_ENCCU |
